# Quantitative proteomics identifies the universally conserved ATPase Ola1p as a positive regulator of heat shock response in *Saccharomyces cerevisiae*

**DOI:** 10.1101/2021.05.12.443772

**Authors:** Stefan Dannenmaier, Christine Desroches Altamirano, Lisa Schüler, Ying Zhang, Johannes Hummel, Silke Oeljeklaus, Hans-Georg Koch, Sabine Rospert, Simon Alberti, Bettina Warscheid

**Affiliations:** Biochemistry and Functional Proteomics, Institute of Biology II, Faculty of Biology, University of Freiburg, 79104 Freiburg, Germany; BIOTEC and CMCB, Technische Universität Dresden, 01307 Dresden, Germany; Institute of Biochemistry and Molecular Biology, ZBMZ, Faculty of Medicine, University of Freiburg, 79104 Freiburg, Germany; Signalling Research Centres BIOSS and CIBSS, University of Freiburg, 79104 Freiburg, Germany

**Keywords:** Proteomics, heat shock protein, protein aggregation, protein misfolding, stress granule, protein self-assembly, Heat shock factor protein 1, translation initiation, protein synthesis

## Abstract

The universally conserved P-loop ATPase Ola1 is implicated in various cellular stress response pathways, as well as in cancer and tumor progression. However, Ola1p functions are divergent between species and the involved mechanisms are only poorly understood. Here, we studied the role of Ola1p in the heat shock response of the yeast *Saccharomyces cerevisiae* using a combination of quantitative and pulse labeling-based proteomics approaches, *in vitro* studies and cell-based assays. Our data show that when heat stress is applied to cells lacking Ola1p, the expression of stress-protective proteins is enhanced. During heat stress Ola1p associates with detergent-resistant protein aggregates and rapidly forms assemblies that localize to stress granules. The assembly of Ola1p was also observed *in vitro* using purified protein and conditions, which resembled those in living cells. We show that loss of Ola1p results in increased protein ubiquitination of detergent-insoluble aggregates recovered from heat-shocked cells. When subsequently cells lacking Ola1p were relieved from heat stress, reinitiation of translation was delayed, whereas, at the same time, *de novo* synthesis of central factors required for protein refolding and the clearance of aggregates was enhanced when compared to wildtype cells. The combined data suggest that upon acute heat stress, Ola1p is involved in the stabilization of misfolded proteins, which become sequestered in cytoplasmic stress granules. This function of Ola1p enables cells to resume translation in a timely manner as soon as heat stress is relieved.

## Introduction

P-loop NTPases play important roles in various cellular processes including protein synthesis, signal transduction, transport and cell division. Ola1 is a member of the highly conserved family of P-loop GTPases and belongs to the subfamily of Obg (*spo<u>0B</u>-associated <u>G</u>TP-binding protein)*-related GTPases (1). In contrast to other members of this class, Ola1 and its bacterial homolog YchF preferentially hydrolyze ATP due to an unusual G4 motif that confers nucleotide binding specificity (2).

Previous studies in bacterial and human cells report a role for Ola1/YchF in different stress response pathways, including the oxidative stress response, integrated stress response and heat shock response (HSR) (3–6). Converging evidence supports a model in which Ola1/YchF acts as a negative regulator of the oxidative stress response (Zhang *et al.*, 2009; Wenk *et al.*, 2012; Hannemann *et al.*, 2016). In contrast, Ola1 was shown to stabilize HSP70 in human cells during heat and oxidative stress conditions (6, 9). Thus, the function of Ola1/YchF in the stress response appears to be divergent and may likely not be explained by a single mode of action. Moreover, information about the function of Ola1 in *S. cerevisiae* (here referred to as Ola1p) is still limited. Global studies revealed that Ola1p interacts with the 26S proteasome (10) and has a potential role in translational fidelity during protein synthesis (11). Recently, Ola1p has been reported to form assemblies upon severe heat shock (12, 13). However, knowledge about Ola1p assembly and its associated function in the HSR is still largely incomplete.

The HSR includes multiple adaption and repair mechanisms that counteract and prevent deleterious effects on cells due to protein destabilization and aggregation (14). Typical features of the HSR include the transcriptional upregulation of heat shock proteins (HSPs) by the transcription factor Hsf1, the inhibition of bulk protein synthesis and increased proteolysis. Cellular fitness during heat shock further depends on other physiological adaptations such as cell wall remodeling and cytoplasmic rearrangements (15). The latter includes the formation of cytoplasmic stress granules (SGs) to promote cellular survival during stress (16). SGs are ribonucleoprotein assemblies of mRNA and proteins that are typically involved in protein synthesis and RNA binding (17). SGs may form via a demixing process known as liquid-liquid phase separation, whereby a homogenous mixture of components (proteins and mRNA) separate from the cytoplasm to assemble into a dense phase, termed condensate or assembly (18). SGs have been reported to regulate translation during stress by the sequestration of translation factors and by storing non-translating mRNAs (19). Moreover, SGs that form during severe heat stress in yeast co-localize with aggregates consisting of ubiquitinated and misfolded proteins. After heat shock, these misfolded proteins are either cleared by proteasomal degradation or refolded by chaperones (20, 21).

Here, we studied the function of Ola1p in the HSR in *S. cerevisiae*. Using quantitative proteomics, we show that deletion of *OLA1* results in the upregulation of a specific set of proteins involved in stress-protective processes, linking Ola1p to the maintenance of cellular proteostatis during heat shock. We found Ola1p in the detergent-resistant pellet fraction of heat-shocked cells. Furthermore, we demonstrate that Ola1p reversibly assembles in a temperature-dependent manner and localizes to cytoplasmic SGs. *In vitro* studies revealed that Ola1p assembly is protein autonomous and coincides with structural rearrangements in the protein. We found that Ola1p reduces protein ubiquitination levels in the detergent-resistant pellet fraction from heat-shocked cells, which indicates that cells may likely be able to faster restore proteostatis upon stress relief. Using pulse labeling experiment, we reveal that global translation reinitiation is markedly reduced in Δ*ola1* cells after heat stress relief. Moreover, deletion of *OLA1* is associated with a specific modulation of the translational program with increased synthesis of chaperones and proteasomal subunits, whose functions are needed to effectively restore a functional proteome for the fast recovery of cells.

## Results

### Deletion of *OLA1* results in the upregulation of stress-protective proteins in cells exposed to heat shock

To investigate the role of Ola1p in the heat stress response in *S. cerevisiae*, we performed a quantitative proteomics study employing native stable isotope labeling by amino acids in cell culture (SILAC) (22). Following metabolic labeling, wildtype and *Δola1* cells were subjected to mild heat stress (42°C, 60 min) and compared to control cells (30°C) by mass spectrometry (MS)-based quantitative proteomics as outlined in **Figure 1a**. More than 100 proteins were significantly increased in abundance in wildtype cells exposed to heat shock, with the strongest increase seen in the levels of heat shock proteins (HSPs) of the Hsp70 (Ssa1, Ssa4), Hsp90 (Hsp82) and Hsp100 (Hsp104) family as well as small HSPs (Hsp26, Hsp42) (**Figure 1b, Supplemental Table S1a**). Notably, expression of Ola1p was not altered upon heat shock, which was confirmed by Western blot analysis (**Figure S1a**). Gene ontology (GO) term enrichment analysis of proteins with increased abundance in heat-stressed cells showed an overrepresentation of proteins involved in cell-protective processes including response to heat and chaperone-mediated protein folding (**Figures 1b** and **S1b, Supplemental Table S1b**). In contrast, factors involved in ribosome biogenesis or RNA processing (e.g. Nsr1, Arx1, Nug1, Mak16) were overrepresented among the proteins with decreased abundance (**Figures 1b** and **S1c, Supplemental Table S1b**).

**Figure 1.**
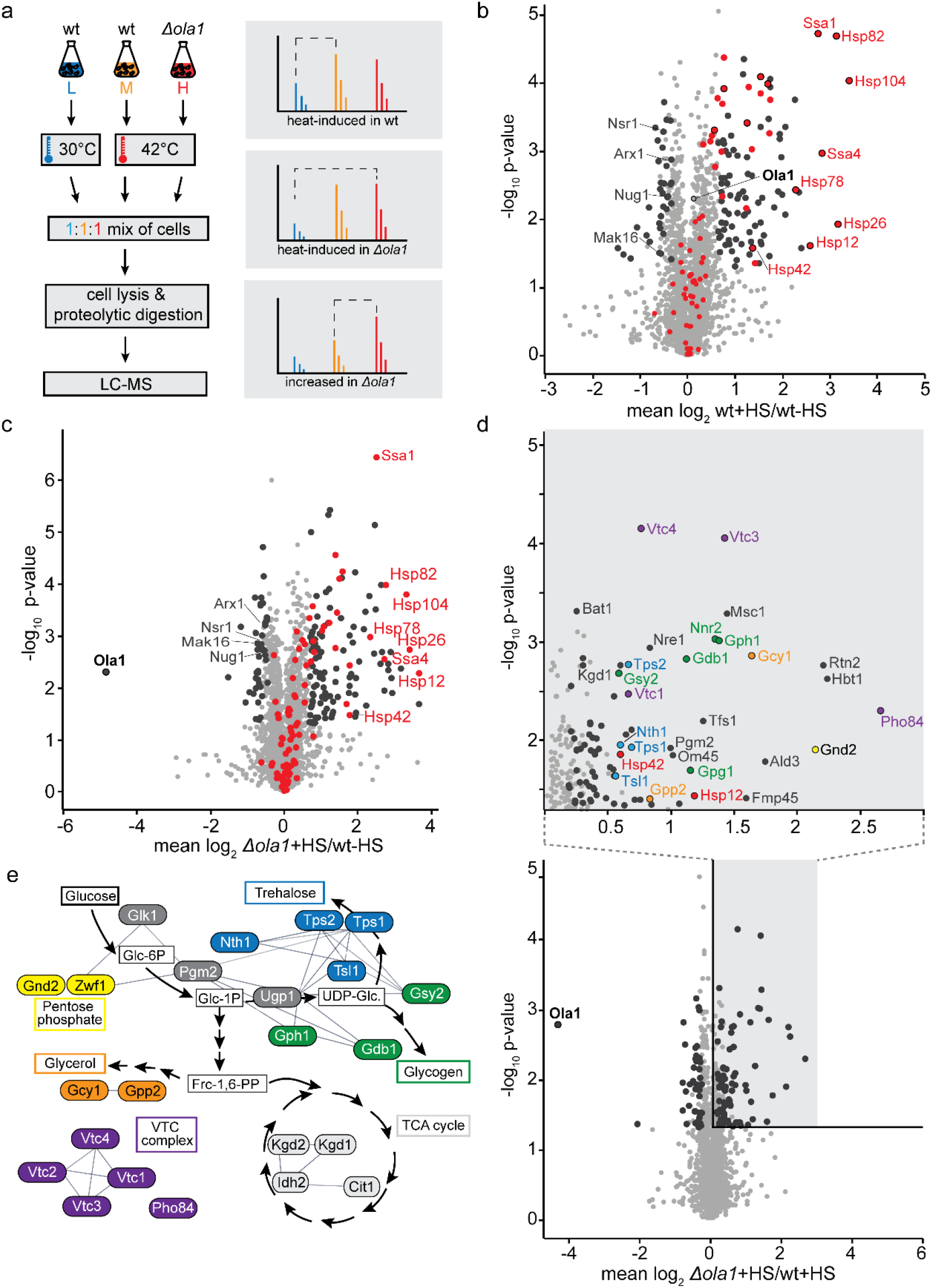
Deletion of *OLA1* results in increased levels of a set of stress-protective proteins during heat shock. a. Triple native SILAC approach employed to study effects of heat stress (42°C, 60 min) on the proteomes of wildtype (wt) and Δ*ola1* cells. b, c. Effect of heat stress (HS) on the proteome of wildtype cells (b) and Δ*ola1* cells (c), analyzed as shown in (a). Larger circles mark proteins significantly changed in abundance following heat stress (i.e., p-value < 0.05 for both t-test and Significance B, n = 4). Proteins annotated with the GO term “Protein Folding” and known chaperones are highlighted in red. d. Bottom, Heat stress-induced changes in the proteome of Δ*ola1* cells *versus* wildtype cell, analyzed as shown in (a). Larger circles mark proteins significantly higher/lower in abundance in Δ*ola1* cells (i.e., p-value < 0.05 for both t-test and Significance B, n = 4). Top, Zoom-in of the shaded area of the volcano plot. Colors correspond to the colors used in (d). e. Affiliation of proteins with significantly higher levels in heat-stressed Δ*ola1* cells to distinct biosynthetic pathways. Lines indicate interactions derived from Cytoscape (67).

We next analyzed the effect of the deletion of *OLA1* in yeast cells exposed to heat stress. As observed in wildtype cells, *Δola1* cells showed a strong induction of HSPs upon exposure to heat stress when compared to unstressed cells (**Figure 1c, Supplemental Table S1a**). Since loss of Ola1p had only a minor effect on the proteome under non-stress condition (**Figure S1d, Supplemental Table S2**), we concluded that the observed changes are specific to heat stress. Moreover, quantitative comparison of the proteome of heat-stressed *Δola1* and heat-stressed wildtype cells revealed that an additional set of stress-associated proteins was increased in abundance (**Figure 1d**). Among these proteins were Hsp12, which helps to maintain membrane organization and prevents protein aggregation (23, 24), the small heat shock protein Hsp42, which suppresses unfolded protein aggregation and sequesters protein aggregates into deposition sites (25, 26), and proteins involved in the synthesis of the chemical chaperone polyphosphate (Vtc1-4). Moreover, our data indicate a rearrangement in energy metabolism as we observed increased levels of proteins involved in the pentose phosphate pathway (Gnd2, Zwf1), TCA cycle (Kgd1, Kgd2, Cit1, Idh2), glycogen synthesis (Gph1, Gdb1, Gsy2), trehalose synthesis (e.g. Tps1, Tsl1) and glycerol synthesis (e.g. Gpp2, Gcy1) (**Figures 1d,** inset, **1e** and **S1e, Supplemental Table S1c**).

Taken together, our data show that loss of Ola1p is counteracted by the upregulation of an additional set of proteins involved in stress-protective processes, suggesting that Ola1p contributes to processes for maintaining proteostasis during heat shock.

### Ola1p is enriched in detergent-insoluble protein aggregates during heat shock

Unbalanced proteostasis, as we observed in *Δola1* cells, is typically associated with the accumulation of protein aggregates. To examine the impact of Ola1p on heat-induced protein aggregation, we performed a triple SILAC approach as outlined in **Figure 1a** and combined it with the enrichment of detergent (NP-40)-insoluble protein aggregates from these cells. In wildtype cells, 185 proteins were found to be enriched (≥ 2-fold; *p-*value < 0.05) in the NP-40 insoluble pellet fraction in response to severe heat stress (46°C, 12 min) (**Figure 2a, Supplemental Table S3a**). Among the 185 detergent-resistant proteins with an increased abundance in the detergent-insoluble pellet fraction of heat-stressed cells, we identified Ola1p and known SG marker proteins such as Ded1, Pab1, and Pub1, as well as numerous chaperones (e.g. Ssb1, Ssb2, Ssz1, Ssa4, Hsp26) (**Figure 2a**). Based on measured MS intensities, cytosolic proteins account for the largest fraction (∼50%) of heat shock-enriched proteins in the pellet fraction, followed by nuclear (27%) and mitochondrial (15%) proteins (**Figure 2b**). Interestingly, proteins of both cytoplasmic SGs and proteasome storage granules, which are assumed to help protect proteasomal components from autophagic degradation (27), are overrepresented in this fraction (**Figure 2c, Supplemental Table S3b**). Our findings underpin an association of misfolded cytosolic proteins and chaperones with cytoplasmic SGs during heat shock conditions (20). Additional computational analysis of protein features revealed that proteins enriched upon heat shock in the NP-40 insoluble pellet fraction tend to be less hydrophobic, have a lower isoelectric point but are on average more abundant and stable compared to the background fraction (i.e. non-enriched proteins) (**Figure 2d, Supplemental Table S3a**).

**Figure 2.**
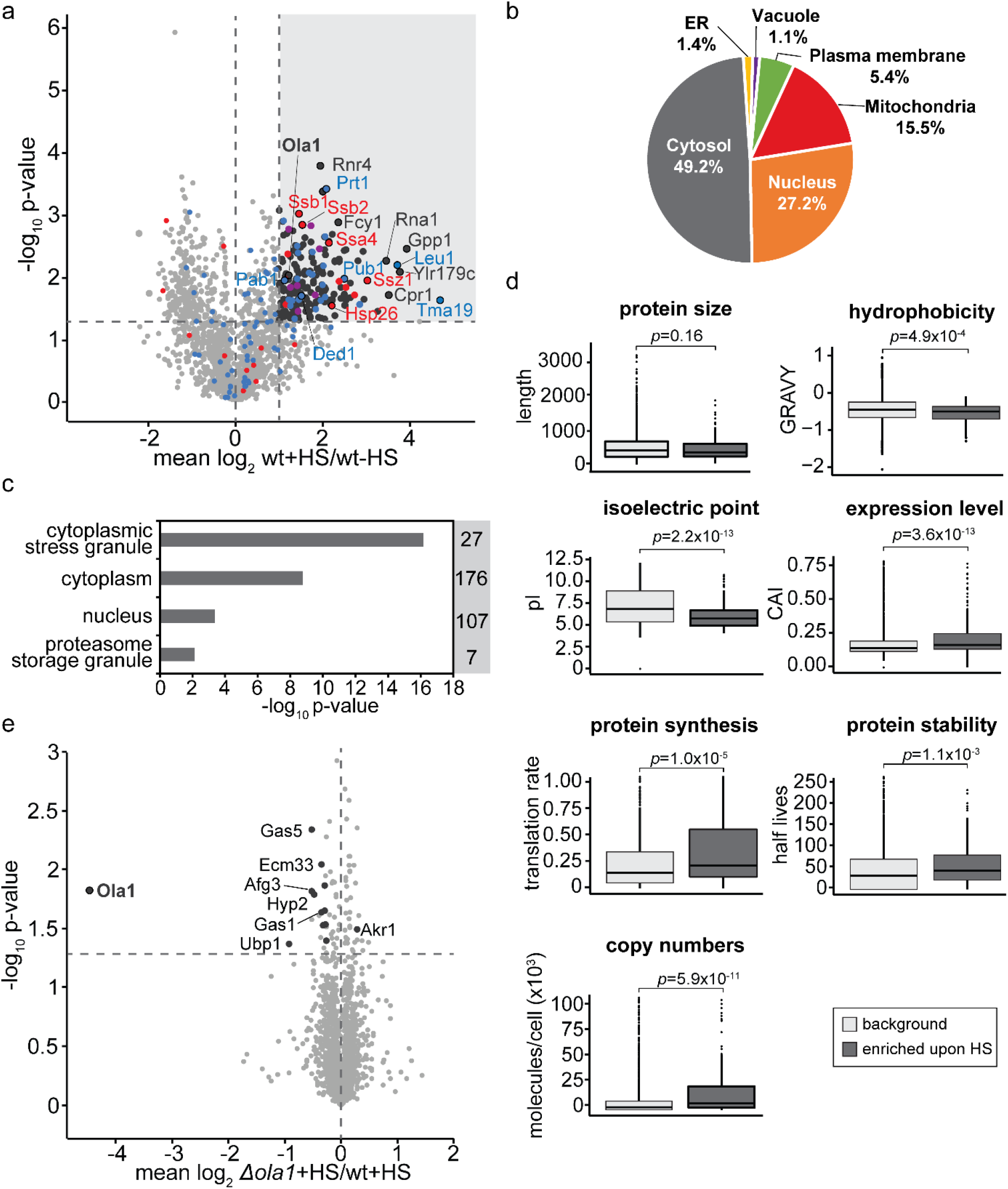
Quantitative Analysis of detergent-insoluble protein aggregates formed during heat shock. a-e. Whole cell extracts of wildtype (wt) and *Δola1* cells were prepared and NP40-insoluble protein aggregates were enriched by centrifugation. Individual fractions of stressed and unstressed cells, taken following severe heat stress (HS; 46°C, 12 min), were analyzed by LC-MS. a, e. The effect of severe heat stress on proteins present in the pellet fraction of wildtype cells (a) and the effect of *OLA1* deletion on proteins in the pellet fraction of heat-stressed cells (e) was assessed by quantitative MS. Larger circles mark proteins significantly altered in abundance (i.e., ≥ 2-fold change and *t-*test *p*-value < 0.05, n = 3). Stress granule proteins are highlighted in blue, chaperones are highlighted in red, proteasome storage granules are highlighted in purple. b, c. Subcellular localization (b) and GO term enrichment analysis (c) of proteins that were at least 2-fold enriched (p-value < 0.05) in the NP-40-insoluble pellet fractions of heat-stressed wildtype cells. Numbers in (c) indicate the number of proteins assigned to the respective term for the domain ‘cellular component’. d. Comparison of the physio-chemical properties of enriched proteins (≥ 2-fold enriched and p-value < 0.05) to the background population (< 2-fold enriched and/or p-value ≥ 0.05). Hydrophobicity and protein expression levels were assessed based on GRAVY scores and the codon adaptation index (CAI), respectively.

As shown in **Figure 2e**, deletion of *OLA1* only had a marginal effect on the protein composition of the detergent-resistant pellet fraction. Among the few proteins with a slightly decreased abundance are the deubiquitinase Ubp1, the translation elongation factor Hyp2 (eIF-5A), and the mitochondrial inner membrane AAA ATPase Afg3, which mediates the degradation of misfolded proteins in mitochondria (28), but has also been shown to be involved in cytoplasmic mRNA translation (29). Thus, while Ola1p localizes to detergent-resistant aggregates formed during heat stress, it has virtually no effect on their protein composition.

### Ola1p assembly is highly sensitive towards heat shock

Based on our finding that Ola1p is present in a detergent-insoluble fraction enriched for protein aggregates that formed during heat stress and has been associated with cytoplasmic SGs (12), we investigated heat-induced changes in the localization of Ola1p in yeast cells expressing C-terminally tagged Ola1p^GFP^ using confocal fluorescence microscopy. We found that cytoplasmic Ola1p^GFP^ assemblies already form at 42°C and strongly increase in number and size when cells were exposed to more severe heat stress (46°C) (**Figure 3a**). Use of the SG marker Pub1^mCherry^ (30) showed that Ola1p^GFP^ co-localizes with heat-induced SGs to a high degree (Pearson’s R-value 0.92) when cells were exposed to severe heat stress (**Figure 3b**). Notably, other stress conditions that typically result in SG formation, such as oxidative stress, did not induce assembly of Ola1p in yeast cells (**Figure S2a**).

**Figure 3.**
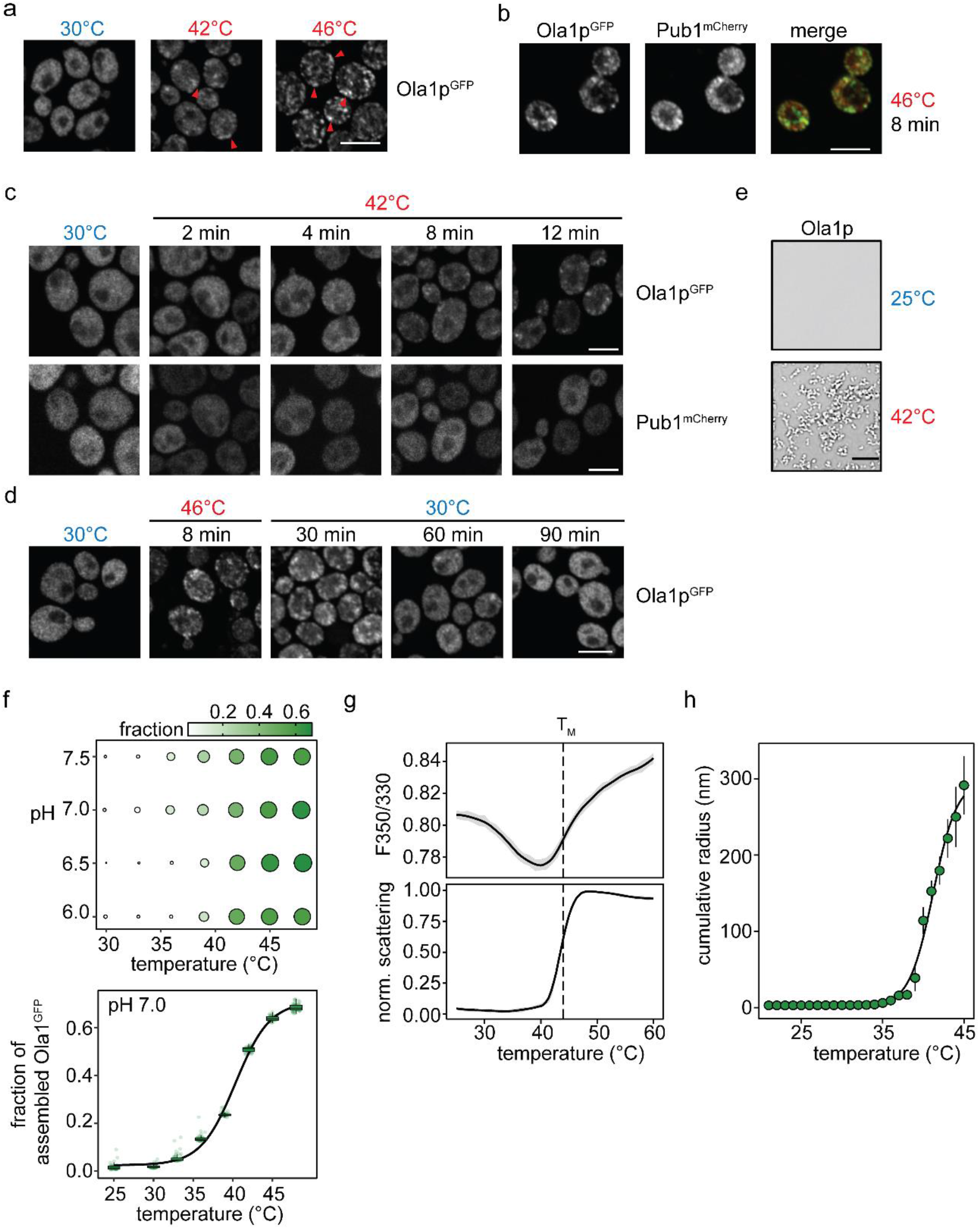
Ola1p assembles upon heat stress and undergoes conformational rearrangement. a. Fluorescence microscopy images of cells expressing endogenous Ola1p^GFP^ exposed to increasing temperatures for 15 minutes. Arrowheads indicate Ola1p^GFP^ assemblies (examples). Scale bar, 5 µm. b, c. Cells expressing endogenous Ola1p^GFP^ were transformed with a plasmid expressing mCherry-tagged Pub1 (Pub1^mCherry^). Co-localization of Ola1p^GFP^ and Pub1^mCherry^ was assessed by fluorescence microscopy after severe heat shock for 8 minutes (b) and at different time points as indicated under mild heat stress conditions (c). Scale bars, 5 µm. d. Ola1p^GFP-^expressing cells were exposed to severe heat shock (46°C; 8 min) followed by recovery at 30°C for 90 minutes and analyzed by fluorescence microscopy at the indicated time points. Scale bar, 5 µm. e. Representative microscopy images of 2.5 µM Ola1p in pH 7.0 buffer before (top) and after heating at 42°C (bottom). Scale bar, 10 µm. f. Heat-induced assembly of 2.5 µM Ola1p (1:10 ratio of Ola1p^GFP^ to Ola1p) after a 10-minute incubation at different temperatures and in buffers of different pH (top). Size and colour of circles reflect the apparent mean fraction of assembled Ola1p^GFP^ calculated from 50 fields of views. The distribution of the apparent fraction of assembled Ola1p^GFP^ in pH 7.0 buffer plotted as a function of temperature is shown on the bottom. g. Triplicate F350/330 (top) and scattering (bottom) measurements of 4 µM Ola1p as a function of temperature. The average transition midpoint (T_M_) is marked with a dashed line. h. Cumulative radius (nm) of 10 µM Ola1p as a function of temperature using DLS. The mean ± SD of three replicates is plotted.

The heat-induced assembly of SG proteins is highly sensitive to the heat stress dosage (severity and time) (16, 31–33). Thus, to gain further insight into the sensitivity of Ola1p to elevated temperatures, cells co-expressing Ola1p^GFP^ and Pub1^mCherry^ were exposed to a milder heat stress (42°C) and imaged in a time course experiment. While Ola1p^GFP^ formed assemblies already after eight minutes at 42°C, Pub1 assembly was not observed at any tested time point (**Figure 3c**). We further tested whether the assembly of Ola1p is reversible. Cells expressing Ola1p^GFP^ were subjected to severe heat stress (46°C) and then shifted to 30°C to monitor the disassembly of Ola1p assemblies over time. While assemblies were still visible at 30 and 60 minutes, Ola1p^GFP^ showed a diffuse distribution throughout the cytosol after 90 minutes of recovery (**Figure 3d**). To conclude, Ola1p rapidly assembles upon heat stress and these assemblies resolve upon stress relief. Moreover, comparison of Ola1p assembly with the SG marker Pub1 shows that Ola1p assembly is more sensitive to heat stress and already assembles at milder heat stresses.

### Ola1p assembles in a temperature-dependent manner *in vitro*

The condensation of proteins has been linked to the presence of intrinsically disordered regions (34). Indeed, many proteins that assemble into SGs have long regions of predicted intrinsic disorder (13). By contrast, Ola1p has a defined three-dimensional structure with no predicted disordered domains (**Figure S2b**). To gain further insight into the heat-induced assembly of Ola1p, we studied its thermostability and assembly properties *in vitro*. Ola1p and Ola1p^GFP^ were purified from insect cells (**Figure S2c**) and studied in buffers of varying pH to mimic the cytosolic pH changes of yeast cells during heat shock. Under non-stress conditions, the cytosolic pH of yeast cells is close to 7.5 and acidifies to values between 6.5 and 7.0 during heat shock (35, 36). Using microscopy, we observed no assembled forms of Ola1p in pH 6.0-7.5 buffers at a concentration of 2.5 µM, which reflects its cellular concentration (for details on the calculation, see the Methods section), whereas it formed assemblies resembling small spherical clusters that grow up to more than 10 µm in size upon increasing the temperature to 42°C (**Figure 3e**). These clusters have gel-like properties and did not dissolve upon decreasing the temperature and increasing the salt concentration (**Figure S2d**). Ola1p assembly was rapid and assemblies already formed after 10 minutes of incubation at 38°C. At temperatures above 42°C, more than half of the Ola1p^GFP^ signal was localized to Ola1p assemblies (**Figure 3f**). Thus, the assembly of Ola1p is highly dependent on temperature and occurs over a temperature and pH range that is typically seen in heat-stressed yeast cells (36).

To determine whether the assembly of Ola1p coincides with a conformational rearrangement in structure, the stability of Ola1p was studied using nano differential scanning fluorimetry (nanoDSF) coupled to light scattering measurements. NanoDSF monitors the intrinsic fluorescence of tryptophan (W) and tyrosine (Y) residues at 330 and 350 nm. The position of W and Y residues in Ola1p are highlighted in **Figure S2b**. Since the fluorescence of W, and in part Y, residues depend on their local environment, conformational rearrangements can be monitored by the analysis of the changes in fluorescence ratio (F350/330) as a function of temperature. Thus, when referring to conformational rearrangements in Ola1p, we report on changes in F350/330. Between 25°C to 38°C, we observed a downward drift in F350/330 as a function of temperature that did not coincide with an increase in scattering from parallel light scattering measurements (**Figure 3g**). Between the temperature range of 38-42°C, however, Ola1p underwent a sharp transition in F350/330 and an increase in light scattering, suggesting Ola1p assembly (**Figure 3g**) as seen in our microscopy data (**Figures 3e** and **3f**). To determine whether the change in F350/330 coincides with Ola1p assembly, we analyzed the transition midpoint (T_M_) of the F350/330 and scattering measurements. At cellular Ola1p concentrations of 2-4 µM, we observed an overlap in the T_M_ of F350/330 and scattering, suggesting that a conformational rearrangement drives Ola1p assembly. At higher concentrations, these events become progressively more uncoupled where the T_M_ of scattering was 1-3°C below the T_M_ of F350/330, suggesting that Ola1p assembly is only marginally concentration-dependent (**Figure S2e**). To gain a better insight into the conformational size of Ola1p, the cumulative hydrodynamic radius of Ola1p was also monitored as a function of the temperature using dynamic light scattering (DLS). At temperatures close to 25°C, Ola1p adopts a monomeric and compact state with a cumulative radius of 3.0 nm (± 0.1), while at temperatures close to the T_M_, the cumulative radius increases steeply (**Figure 3h**). To conclude, Ola1p assembly is highly dependent on an increase in temperature *in vitro*. In yeast cells, Ola1p forms visible assemblies at 42°C and we show that Ola1p heat-induced assembly is protein autonomous and coincides with a conformational rearrangement at similar temperatures, pH and concentrations *in vitro*. Moreover, we found that Ola1p adopts a compact state at non-stress temperatures and rapidly assembles at elevated temperatures.

### Ola1p does not affect the activation of the HSR by Hsf1

Yeast cells need to properly sense temperature changes to activate the HSR. Our data show that Ola1p has the intrinsic property to rapidly assemble upon an increase in temperature. We hypothesized that this could contribute to an early and efficient activation of the HSR. To address this, we monitored the phosphorylation status of Hsf1 as a measure for HSR activation using phos-tag gel analysis. In both wildtype and *Δola1* cells, Hsf1 migrated as a double band on phos-tag gels at 30°C and shifted to a more highly phosphorylated species with increasing temperatures (**Figure 4a**). A time course experiment showed that Hsf1 gradually shifted to more highly phosphorylated species with prolonged incubation at 42°C in both wildtype and *Δola1* cells (**Figure 4b**). However, based on this phosphorylation analysis, we did not observe considerable differences in Hsf1 activation between wildtype and *Δola1* cells.

**Figure 4:**
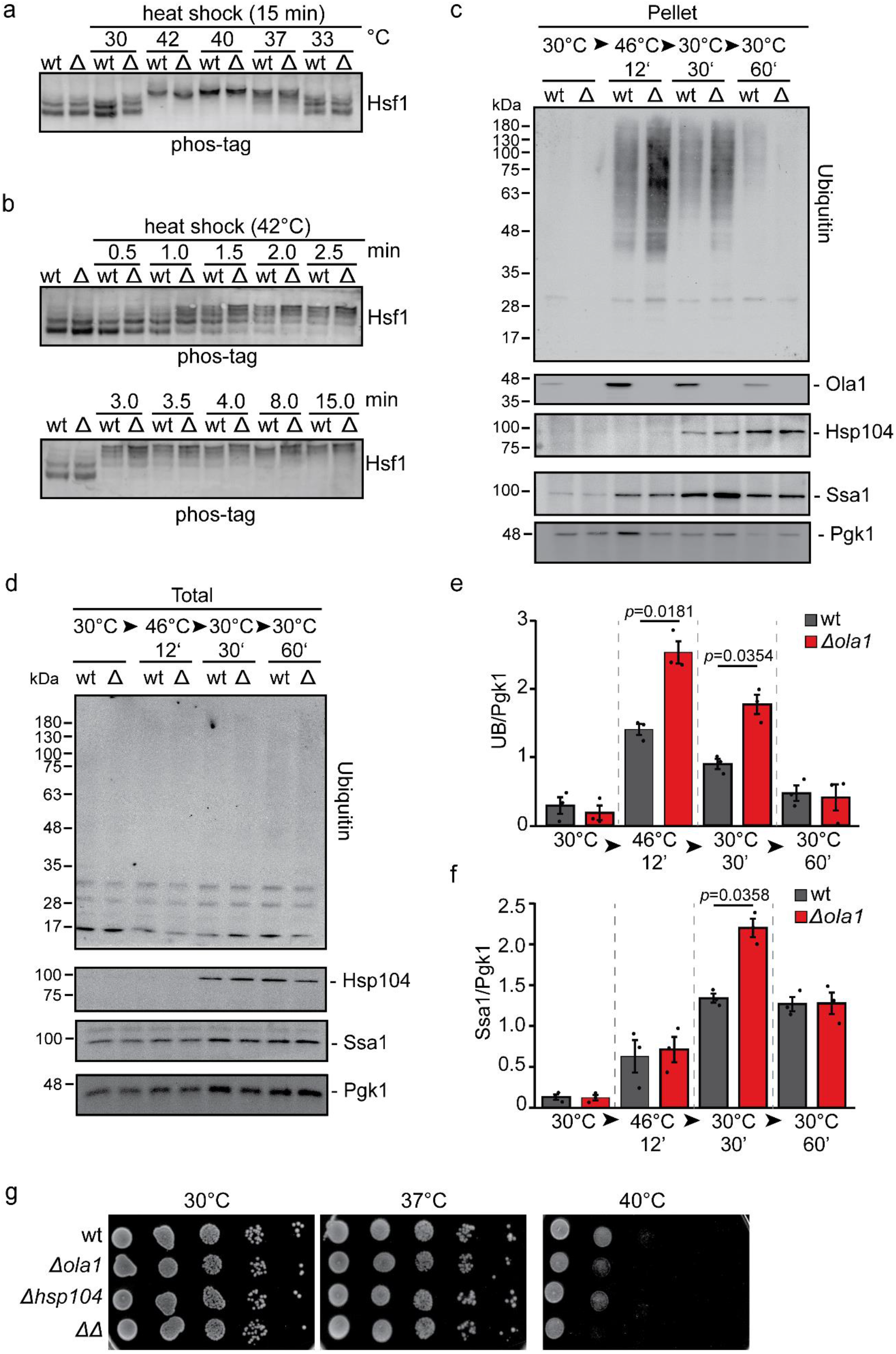
Deletion of *OLA1* does not affect Hsf1 phosphorylation but causes increased ubiquitination of detergent-resistant protein aggregates. a, b. Wildtype (wt) and Δ*ola1* (Δ) cells were subjected to heat shock at different temperatures (a) or times (b) as indicated. Whole cell extracts were prepared, resolved on phos-tag gels and analyzed via immunoblotting with α-Hsf1. c-f. Whole cell extracts (Total) of wildtype (wt) and Δola1 cells (Δ) were prepared and NP40-insoluble protein aggregates (Pellet) were enriched by centrifugation. Individual fractions of stressed and unstressed cells, taken following severe heat stress (46°C, 12 min) and during recovery (30°C) from heat shock, were analyzed by immunoblotting. Levels of protein ubiquitination (e) and Ssa1 (f) in protein aggregates (c) and whole cell extracts (d) were analyzed by immunoblotting. Shown are representative immunoblots of three independent biological replicates. Signals for ubiquitin (UB) and Ssa1 in pellet fractions were quantified and normalized to the signal of the loading control Pgk1 in whole cell extracts (c, d). Error bars indicate SD. g. Serial dilutions of wildtype (wt) and mutant cells (ΔΔ: *Δola1Δhsp104*) were spotted onto YPD plates and incubated for 2 days at 30°C and 37°C or 4 days at 40°C.

### Loss of Ola1p results in higher ubiquitination levels in protein aggregates

So far, we showed that Ola1p localizes to SGs and enriches with detergent-insoluble protein aggregates upon heat shock and deletion of *OLA1* results in increased levels of specific stress-protective proteins including the cytosolic molecular chaperone Hsp42, which is crucial for protein homeostasis in *S. cerevisiae* (25, 26). However, Ola1p does neither affect the composition of the insoluble proteome nor the activation of Hsf1. To further check whether Ola1p has a potential impact on protein misfolding, we next analyzed global ubiquitination levels indicative for protein misfolding (37). We monitored changes in global protein ubiquitination levels in the detergent-resistant pellet fraction of wildtype *versus Δola1* cells by immunoblot analysis (**Figure 4c**). As expected, protein ubiquitination levels were markedly increased in the detergent-resistant pellet fractions of heat-stressed cells (**Figures 4c**). For quantification, the immunoblot signals detected in the detergent-resistant pellet fractions were normalized to the signals of Pgk1 from whole cell lysates (**Figure 4d**). As a result, we determined significantly increased ubiquitination levels in *Δola1* cells during severe heat stress (46°C, 12 min) compared to wildtype cells (**Figure 4e**). During the recovery from heat stress, ubiquitination levels gradually decreased in wildtype and *Δola1* cells. We observed a similar decrease in Ola1p levels in the pellet fraction during recovery (**Figures 4c**). Notably, overexpression of Ola1p from a plasmid in heat-shocked *Δola1* cells partially restored ubiquitination levels back to wildtype levels (**Figure S3a**).

Previous work showed that protein aggregates are cleared in an Ssa1-Hsp104-dependent manner (20, 38). Thus, we next monitored the levels of Ssa1 and Hsp104 during heat stress and stress relief. Both Ssa1 and Hsp104 were present in considerably larger amounts in the pellet fraction of *Δola1* cells after 30 minutes of recovery (**Figures 4c, 4f** and **S3b**), despite equal expression in *Δola1* and wildtype cells (**Figure 4d**; see also **Supplemental Table S1**). Interestingly, we also identified a genetic interaction between *OLA1* and *HSP104*. In plating experiments, we observed a temperature-sensitive growth phenotype at 40°C in *OLA1* and *HSP104* double knockouts, which was less pronounced in the respective single mutants (**Figure 4g**).

To conclude, increased levels of Ssa1 and Hsp104 in the detergent-resistant pellet fraction of *Δola1* cells and the growth defect of *Δola1Δhsp104* cells seen under heat stress suggest that Ola1p is likely involved in the stabilization of misfolded proteins during heat shock, thereby allowing cells to faster restore a functional proteome upon stress relief.

### Ola1p promotes efficient reinitiation of translation after stress relief

In response to heat shock, cells attenuate global translation to prevent further protein damage. Upon stress relief, a functional proteome needs to be restored and translation effectively reinitiated to allow cells to recover (20). Based on our finding that protein ubiquitination in the detergent-resistant pellet fraction is increased in *Δola1* cells during heat stress, we hypothesized that Ola1p has a stabilizing effect on such proteins. In this case, we would also expect that reinitiation of global translation is less effective in *Δola1* compared to wildtype cells after heat stress relief. To address this, we performed *in vivo* pulse labeling experiments and followed the incorporation of ^35^S-methionine/-cysteine into nascent proteins of wildtype and *Δola1* cells during recovery from heat stress (42°C, 10 min). Autoradiographs of radiolabeled proteins show an increase in signal intensities with time, indicating that cells are able to reinitiate translation after stress relief (**Figure 5a**). However, translation reinitiation was significantly reduced in cells deleted for *OLA1* compared to wildtype cells after heat stress (**Figure 5b**), which was not observed under non-stress condition (**Figures S4a** and **S4b**). We thus conclude that reinitation of translation after heat shock is delayed in cells lacking Ola1p. Based on this finding, we next analyzed changes in the synthesis rate of proteins in wildtype and *Δola1* cells during the recovery from heat stress using a pulsed SILAC (pSILAC) approach (39) and quantitative MS (**Figure 6a**). With this approach, we distinguished newly synthesized proteins from pre-existing ones and followed their synthesis between 30 and 120 minutes after heat stress (42°C, 30 min). The global relative translation levels of wildtype and *Δola1* cells were determined at 30, 60, 90 and 120 min of recovery based on the overall distribution of the SILAC ratio H/M (i.e. ratio *Δola1*/wt) (40) (**Figure 6b, Supplemental Table S4a**). At early timepoints of heat stress recovery (i.e. 30 and 60 min), the ratio *Δola1*/wt was below 1, indicating that protein synthesis during recovery from heat stress is reduced in *Δola1* cells in line with our radioactive pulse labeling analysis (see **Figure 5**). We next determined the changes in the synthesis of individual proteins from wildtype *versus Δola1* cells upon recovery from heat shock (**Figures 6c** and **6d**, **Supplemental Table S4a**). Hierarchical cluster analysis identified three main protein clusters. 609 proteins showed a reduced synthesis at early timepoints of recovery (cluster 1), while the synthesis of 120 proteins (cluster 2) was increased upon deletion of *OLA1*. For 594 proteins (cluster 3), no effect on synthesis during heat stress recovery was observed (**Figures 6c** and **6d**). Relative synthesis profiles for representative proteins of clusters 1-3 are depicted in **Figure 6e** and **Figure S5a**. GO term enrichment analysis revealed that components of the preribosome and ribosome as well as proteins involved in translation initiation and RNA localization were overrepresented in cluster 1 (**Figure 6f**, **Supplemental Table S4b**). Among the proteins overrepresented in cluster 2 were many chaperones including Hsp12, Hsp26, Hsp60, Hsp78, Hsp82, Hsp104 and Ssa1 as well as proteins involved in amino acid biosynthesis and components of the proteasome. In contrast, proteins of cytosolic SGs, proteasome storage granules or intracellular organelles, as well as factors involved in nucleotide synthesis and protein transport were overrepresented in cluster 3 (**Figure S5b**). Notably, when we overexpressed Ola1p from a plasmid in *Δola1* cells, we could rescue the observed phenotypes (**Figure 6g, Supplemental Table S5**), underscoring that these cellular responses directly relate to Ola1p function. In summary, our data show that Ola1p promotes the efficient reinitiation of translation after heat stress. Although global translation was reduced in *Δola1* cells, the synthesis of numerous factors of the proteostasis network was considerably increased. Higher synthesis levels of chaperones and proteasomal subunits in *Δola1* cells support our notion that Ola1p reduces damage of aggregation-prone proteins during heat stress and thereby in turn aids in the efficient reinitation of global translation after stress relief.

**Figure 5.**
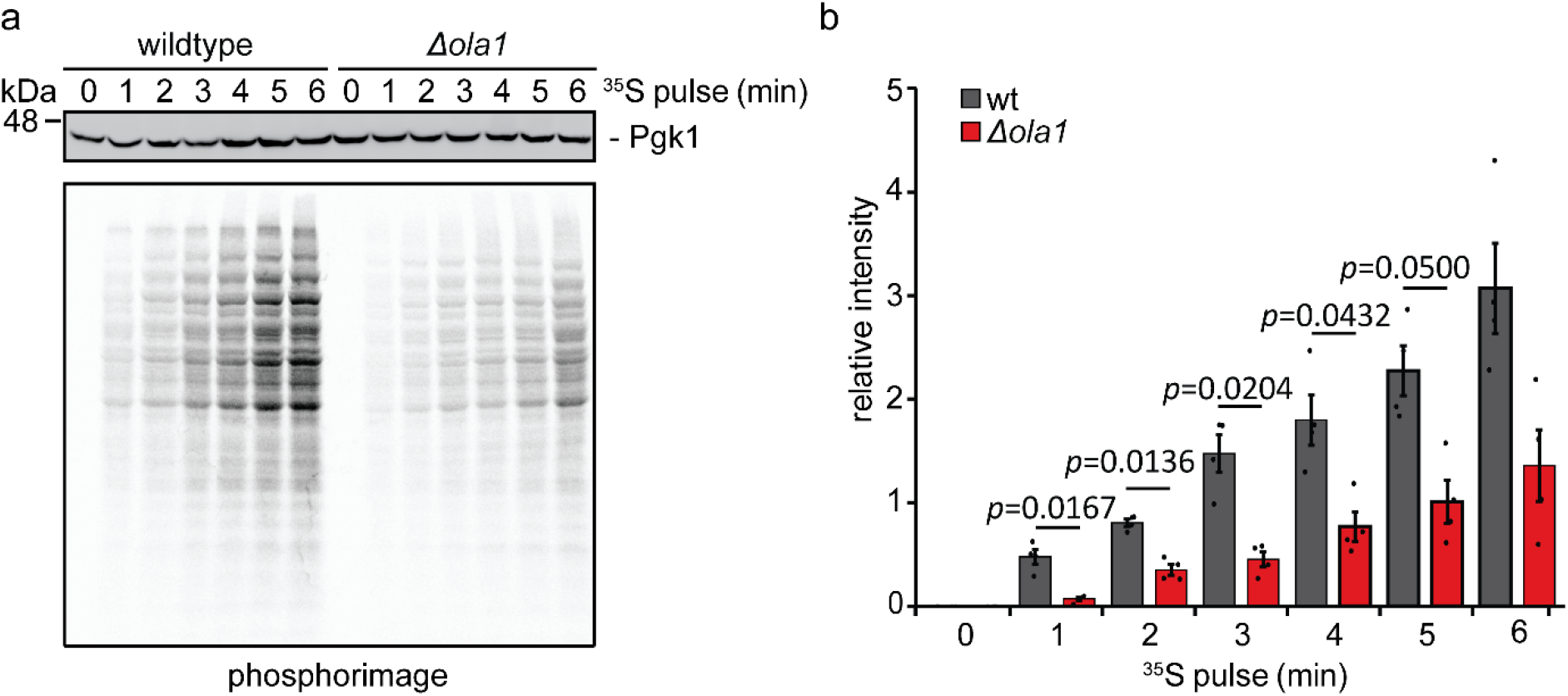
Reinitiation of translation after stress relief is impaired in *Δola1* cells. a, b. Wildtype (wt) and *Δola1* cells in logarithmic growth phase were subjected to heat stress at 42°C for 10 minutes. After addition of ^35^S-methionine/-cysteine, cells were transferred to 30°C, and incorporation of ^35^S into newly synthesized proteins was monitored by autoradiography at the indicated timepoints (a). Autoradiographic intensities (n = 4) were quantified and normalized to the loading control Pgk1 (b). Error bars indicate SD.

**Figure 6.**
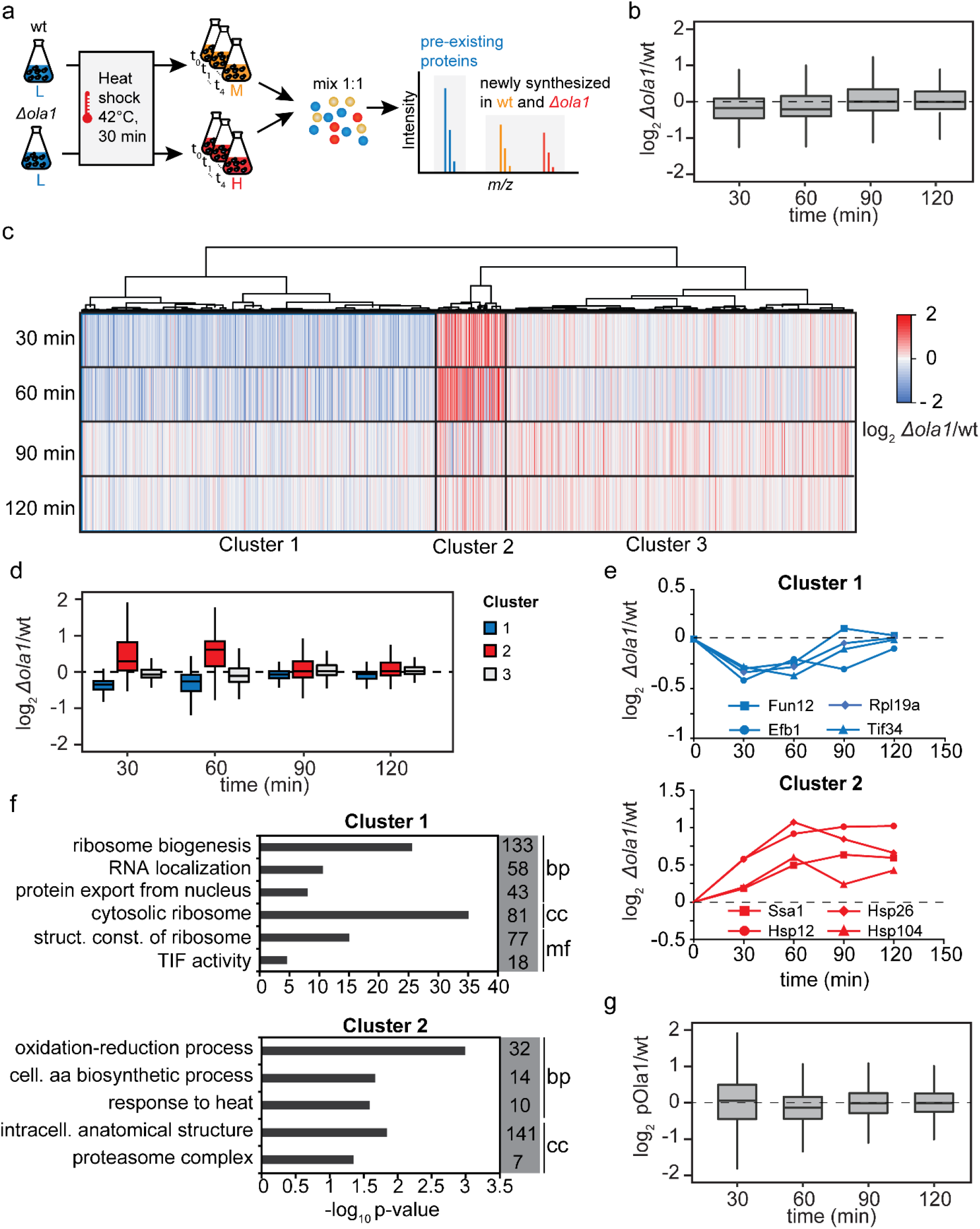
Cells lacking Ola1p specifically modulate their translational program to restore a functional proteome after heat stress relief. a. Pulsed SILAC-based proteomics approach used to study protein synthesis in wildtype (wt) and *Δola1* cells during recovery from heat shock (42°C, 30 min, n = 3). b. Relative differences in global protein synthesis between wildtype and *Δola1* cells 30, 60, 90 and 120 minutes after heat shock. c, d. Hierarchical cluster analysis of *Δola1*/wt ratios reflecting the difference in protein synthesis between wildtype and *Δola1* cells. e. Temporal profiles of relative protein synthesis of proteins representative for clusters 1 and 2 as determined by hierarchical cluster analysis. f. GO term overrepresentation analysis of clusters 1 and 2 for the domains ‘cellular component’ (cc), ‘molecular function’ (mf), and ‘biological process’ (bp). Listed are selected terms overrepresented in a given cluster and number of proteins assigned to each term that are present in the respective cluster. struct. const., structural constituent; TIF, translation initiation factor; (intra)cell., (intra)cellular; aa, amino acid. g. Relative differences in global proteins synthesis between wildtype (wt) and *Δola1* cells overexpressing Ola1p (pOla1p) 30, 60, 90 and 120 minutes after heat shock (42°C, 30 min; n = 3). Data were obtained using a pulsed SILAC approach as described in (a).

## Discussion

Different roles of Ola1/YchF in cellular stress response pathways have been reported (3, 5, 6, 41). Furthermore, *OLA1* is overexpressed in various cancers (42, 43), which underscores the interest in better understanding Ola1/YchF functions. Previous work showed that human OLA1 stabilizes HSP70 during heat shock (6, 9). However, characterization of Ola1/YchF functions is challenging due to its different modes of action, and thus poses a need for systematic studies. To address this, we employed quantitative and functional proteomics methodology to globally analyze the role of Ola1p in maintaining a functional proteome in *S. cerevisiae* cells during heat stress. Our quantitative proteome data revealed a specific role of Ola1p during heat stress. The deletion of *OLA1* led to the increased expression of the molecular chaperones Hsp12 and Hsp42, as well as proteins involved in the synthesis of stress-protective molecules such as polyphosphate and specific metabolic pathways (e.g. for the synthesis of glycogen, glycerol, and trehalose) during heat stress. Hsp42 has been described as a sequestrase that coaggregates with misfolded proteins and sequesters them to distinct cellular sites. Thereby, the properties of misfolded proteins are altered in a way that facilitates their disaggregation by Ssa1/Hsp104 (25, 44). Hsp12 is an intrinsically unstructured protein with a cytosolic pool that prevents protein aggregation and a membrane-associated pool with a potential role in membrane remodeling (23, 24). In addition, a synergetic effect between Hsp12 and trehalose in stabilizing the activity and preventing the aggregation of proteins has been recently reported (24). Polyphosphate has been identified to function as “chemical chaperone” for protecting proteins from unfolding and aggregation under stress conditions (45). Thus, our data reveal that cells compensate for the loss of Ola1p by expressing stress-protective molecules during heat shock and, moreover, suggest a potential link between Ola1p and protein misfolding.

To examine a functional connection between Ola1p and protein misfolding, we performed a quantitative analysis of detergent-insoluble protein aggregates formed during heat shock. This detergent-resistant pellet fraction is mainly composed of highly expressed cytosolic, nuclear and mitochondrial proteins. Notably, we identified Ola1p, chaperones (Ssb1, Ssb2, Ssz1, Ssa4, Hsp26) and other known SG components to be enriched in this fraction during heat stress, which is in line with previous work defining heat-induced SGs as mixed assemblies of SG components and protein aggregates (20). Deletion of *OLA1* only had a minor impact on the formation and the composition of these detergent-resistant protein aggregates, which has also been reported for other “positive regulators” of the HSR like the chaperones Hsp26, Hsp70 or Hsp104 (20). Considering that we identified increased levels of the sequestrase Hsp42 in *Δola1* cells upon heat stress, it appears conceivable that higher levels of Hsp42 may in part compensate for the deletion of *OLA1* and explain why only a minor effect was seen on the formation and composition of the detergent-resistant protein pellets during heat stress.

In line with previous work classifying Ola1p as a “superaggregating” protein (12), we found that Ola1p assemblies form rapidly even at mild heat stress and this heat stress-specific assembly is reversible after stress relief. Ola1p assembly was also characterized *in vitro*. Ola1p was diffuse under non-stress conditions and formed gel-like clusters in a temperature and pH range that corresponds to the conditions seen in the cytosol of heat-stressed yeast cells. We found that Ola1p assembly is protein autonomous and coincides with a conformational rearrangement in protein structure over a concentration range seen in cells. *In vitro*, Ola1p rapidly assembles at temperatures above 40°C, a similar temperature at which another SG protein, the translation factor Ded1p, also undergoes a conformational change and assembles into higher-order structures (33). In this case, the assembly of Ded1p was identified as an integral part of an extended HSR program to aid in the production of stress-protective proteins while repressing the production of housekeeping proteins. Due to its high sensitivity and fast assembly upon elevated temperatures, we further speculated that Ola1p may potentially be involved in the efficient activation of the HSR. However, based on our analysis of Hsf1 phosphorylation during heat shock, which is a hallmark of HSR activation, we could not obtain evidence for a role of Ola1p in HSR activation by Hsf1.

Cells activate the HSR to overcome phases of stress by two arms. First, by reducing heat shock-induced damage of proteins. Second, by enabling the efficient restart of translation upon stress relief. As a measure for heat-induced damage, we analyzed ubiquitination levels of proteins in the detergent-resistant pellet fractions from heat-shocked cells. In the absence of Ola1p, ubiquitination levels were increased in heat-induced protein aggregates, which could be reversed by re-expression of Ola1p from a plasmid. In line with these findings, levels of Hsp104/Ssa1 in the detergent-resistant pellet fraction of *Δola1* cells were increased during the recovery from heat shock. Moreover, cells with a deletion of *OLA1* and *HSP104* are more sensitive to heat stress compared to wildtype and singe deletion strains. Taking these data into account, we put forward the notion that Ola1p associates with misfolding proteins and may likely mitigate unfavorable thermally induced conformational changes to prevent further damage to proteins. Recently published fluorescence recovery after photobleaching experiments have revealed that Ola1p exhibits an extremely low mobility after heat-induced assembly (13), which speaks in favor for a direct association with immobile protein aggregates.

Compelling evidence suggests that the coordinated release of preinitiation complexes and unfolded proteins from heat-induced SGs fine-tunes reinitiation of translation once these SGs are disassembled by Hsp104/Ssa1 chaperones and the protein quality control system has cleared the cytosol from protein aggregates. Notably, translation reinitiation during recovery from stress is tunable and corresponds to the proteostatic state of the cell (20). We therefore anticipated that differences in the level of protein damage should affect translation reinitation after stress relief. Indeed, in the absence of Ola1p cells fail to efficiently resume translation during the recovery from heat shock. Most importantly, time-resolved analysis of the synthesis profiles of hundreds of individual proteins after heat stress relief revealed that deletion of *OLA1* affects translation reinitiation of some, but not all, newly synthesized proteins. While the synthesis of ribosomal proteins and translation initiation factors was reduced up to 2 hours upon stress relief, numerous chaperones including the central factors Hsp104, Ssa1, Hsp26 and Hsp12 and subunits of the proteasome required for the efficient clearance of protein aggregates were synthesized in higher amounts in *Δola1* cells during the recovery from heat stress.

In summary, we have identified a protective role of Ola1p in yeast cells during heat shock. Our data suggest a function of Ola1p in mitigating protein misfolding and damage during heat stress, which eventually enables cells to efficiently reinitate global translation after heat stress relief. Such a role of Ola1p is further supported by its highly temperature-sensitive and rapid assembly, its reversible localization to heat-induced SGs and its constitutive expression at relatively high copy numbers (54,859 copies per cell (46)). Ola1p may therefore qualify as an “early responder” of the HSR to alleviate damage to misfolding proteins. However, it remains to be elucidated how exactly Ola1p stabilizes aggregation-prone proteins during heat stress and whether this mode of action is conserved from yeast to humans.

## Experimental procedures

### Plasmids and cloning techniques

Plasmids used in this study are listed in Table 1. Plasmids were amplified using the *Escherichia coli* strain DH5α and transformed into competent yeast cells following standard procedures. The strains Ssa1^GFP^*Δola1* and *Δola1Δhsp104* were generated by gene deletion via homologous recombination as described before (47). The kanMX4 cassette was amplified from gDNA of Δ*ola1* cells using the Δ*ola1* forward and reverse primers (Table ***2***) and transformed into BY4741 Ssa1^GFP^ cells. The His3MX cassette was amplified from pFA6a using the the Δ*hsp104* forward and reverse primers (Table ***2***) and transformed into BY4741 *Δola1* cells.

**Table 1:** Plasmids used in this study.

| Name | Description | Internal ID |
| --- | --- | --- |
| pOla1p-FLAG | pESC-URA-Ola1-FLAG | P142 (this study) |
| pPub1-mCherry | pRP1362/pRP1363-LEU-Pub1-mCherry | pRP1661 (64) |
| pFA6a | pFA6a-GFP(S65T)-HIS3MX6 | P555 (47) |
| pOCC102-Ola1 | pOEM1-His6-MPB-Ola1 | L-621 (this study) |
| pOCC120-Ola1-monoGFP | pOEM1-MPB-Ola1-monoGFP-His6 | L-571 (this study) |

**Table 2:**
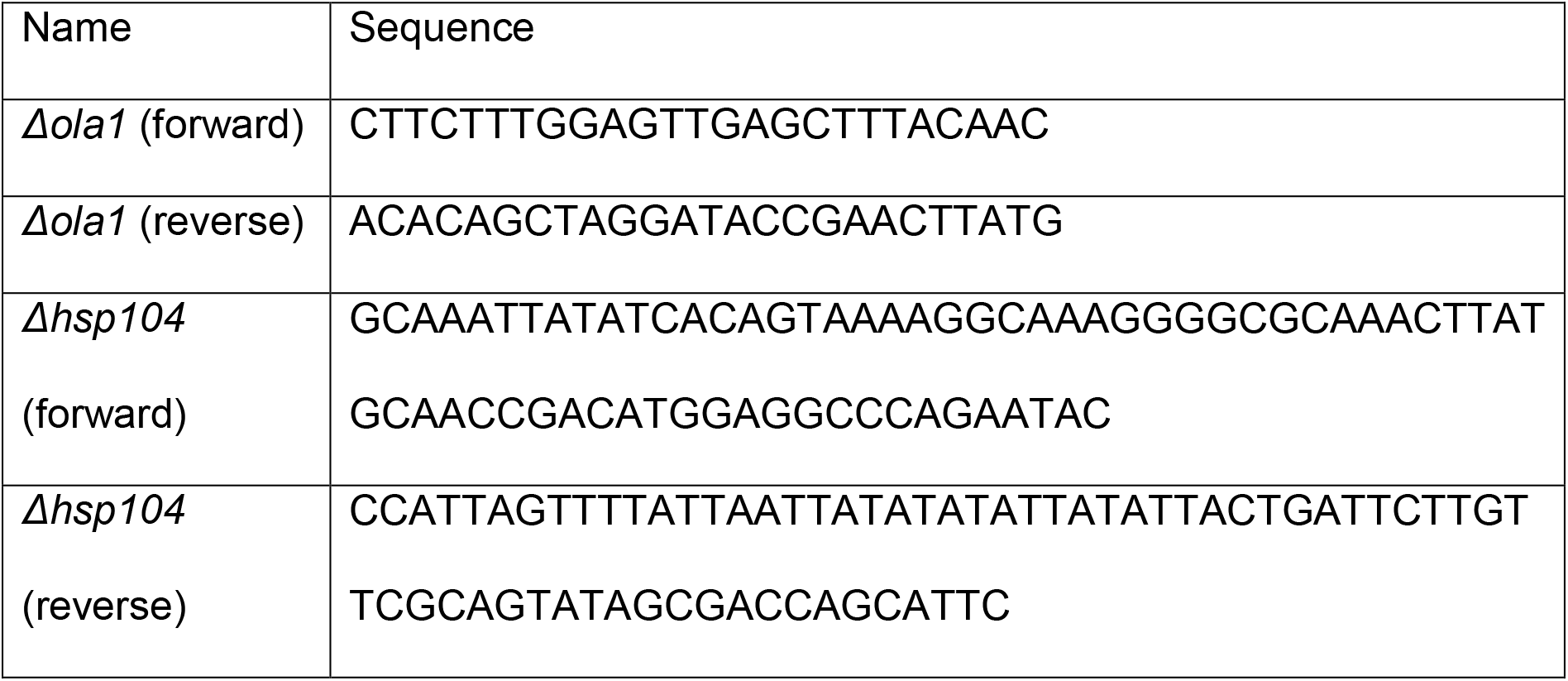
Oligonucleotides used in this study.

### Cultivation of yeast cells and metabolic labeling

*S. cerevisiae* strains used in this study were BY4741 and derivatives thereof (48). Yeast strains are listed in Table 3**Table *3***. Cells were cultivated in liquid medium at 30°C and 160 rpm and harvested in exponential growth phase (OD 0.5 - 1.5) by centrifugation (2 min at 1,600 x g and 4°C), unless otherwise noted.

**Table 3:** Yeast strains used in this study.

| Strain | Description | Genotype | Source |
| --- | --- | --- | --- |
| Wildtype (wt) | BY4741 | *MATa; ura3Δ0, leu2Δ0, his3Δ1, met15Δ0* | Euroscarf |
| *Δola1* | Deletion of *Ybr025c* in BY4741 cells | *MATa; ura3Δ0, leu2Δ0, his3Δ1, met15Δ0, ybr025c:kanMX4* | Euroscarf |
| *Δhsp104* | Deletion of *Yll026w* in BY4741 cells | *MATa; ura3Δ0, leu2Δ0, his3Δ1, met15Δ0, yll26w:kanMX4* | Euroscarf |
| *Δola1*<br>*Δhsp104* | Deletion of *Yll026w* in BY4741 *Δola1/* cells | *MATa; ura3Δ0, leu2Δ0, his3Δ1, met15Δ0, ybr025c:kanMX4, yll026w:his3MX6* | This study |
| *Ola1*<sup>GFP</sup> | BY4741, *Ybr025c*<sup>GFP</sup> (chromosomal) | *MATa; ura3Δ0, leu2Δ0, his3Δ1, met15Δ0, YBR025c:GFP-His3MX* | Euroscarf |
| *Ssa1*<sup>GFP</sup> | BY4741, *Yal005c*<sup>GFP</sup> (chromosomal) | *MATa; ura3Δ0, leu2Δ0, his3Δ1, met15Δ0, Yal005c:GFP-His3MX* | Euroscarf |
| *Ssa1*<sup>GFP</sup><br>*Δola1* | BY4741, *Yal005c*<sup>GFP</sup> (chromosomal),<br>Deletion of *Ybr025c* | *MATa; ura3Δ0, leu2Δ0, his3Δ1, met15Δ0, Yal005c:GFP-His3MX, ybr025c::kanMX4* | This study |

For fluorescence microscopy, enrichment of detergent-insoluble protein aggregates and ^35^S pulse labeling experiments, cells were grown in synthetic complete (SC) medium (0.17% [w/v] yeast nitrogen base without amino acids, 0.5% [w/v] ammonium sulfate, 2% [w/v] glucose, pH adjusted to 5.5) supplemented with appropriate amino acids (49). For analysis of Ola1p expression and growth tests, yeast cells were cultured in YPD medium (1% [w/v] yeast extract, 2% [w/v] peptone, 2% [w/v] glucose) or on YPD plates (YPD medium supplemented with 2% [w/v] agar).

MS-based loss-of-function studies and pSILAC experiments were performed following the 2nSILAC strategy (22). In brief, cells were grown in SC medium supplemented with adenine, L-histidine, L-leucine, L-methionine, L-tryptophane, uracil (20 mg/l each), L-arginine, L-lysine (50 mg/l each) and L-proline (200 mg/l). Precultures with a starting OD_600_ of 0.1 were incubated until they reached an OD_600_ of 3-4 and used to inoculate main cultures with an initial OD_600_ of 0.025. Metabolic labeling was performed using stable isotope-coded “heavy” arginine (13C6/15N4; Arg10) and lysine (13C6/15N2; Lys8) or “medium-heavy” arginine (13C6/14N2; Arg6) and lysine (2H4; Lys4) instead of the corresponding “light” counterparts.

To induce heat stress, yeast cells were harvested in logarithmic growth phase (OD_600_ 0.5-1.5) by centrifugation (2 min at 1,600 x g and RT), resuspended in the appropriate pre-warmed medium and incubated at 42°C or 46°C (as indicated) in a water bath shaker. After application of heat stress, cells were harvested by centrifugation (2 min at 500 x g and 4°C) and shock-frozen in liquid nitrogen.

### Enrichment of detergent-insoluble protein aggregates

Cells (200-250 OD_600_ units) were resuspended in 2 ml of DTT buffer (100 mM Tris/H_2_SO_4_ pH 9.4, 10 mM DTT), equally divided into two reaction tubes and incubated for 10 min at 1,000 rpm and 30°C. Cells were harvested by centrifugation (5 min, 1,500 x g, 4°C) and washed with 1 ml of Zymolyase buffer (50 mM potassium phosphate pH 7.4, 1.2 M sorbitol). Cells were again resuspended in 1 ml of Zymolyase buffer, 4 mg/ml Zymolyase 20-T (MP Biomedicals Life Sciences) were added, and samples were incubated for 20 min at 1,000 rpm and 30°C. After centrifugation for 5 min at 900 x g and 4°C, 300 mg glass beads were added to each reaction tube and cells were lysed by two cycles of bead beating using a Minilys® homogenizer for 3 min at 3,000 rpm. Between the cycles, samples were cooled on ice for 3 min. Cell debris was removed in two centrifugation steps (5 min, 5,000 x g, 4°C) and supernatants were pooled, followed by centrifugation for 20 min at 20,000 x g and 4°C. To solubilize membrane proteins, the pellet was resuspended in 320 µl of aggregate buffer (50 mM potassium phosphate pH 7.0, 1.2 M sorbitol, 1 mM EDTA, 1 mM PMSF, 8 µM antipain-HCl, 0.3 µM aprotinin, 1 mM bestatin-HCl, 10 µM chymostatin, 5 µM leupeptin-H_2_SO_4_, 1.5 µM pepstatin A) and 80 µl of 10% (v/v) NP-40 by pipetting and vortexing. Aggregates were collected by centrifugation (20 min, 20,000 x g, 4°C), the pellet was resuspended as described above, followed by another centrifugation step for 20 min at 20,000 x g and 4°C. Detergent-insoluble protein aggregates were resuspended in sample buffer (0.3 M Tris/HCl pH 6.8, 2% [v/v] SDS, 10% [v/v] glycerol, 100 mM DTT, 6 mM bromophenol blue) and incubated for 10 min at 1,100 rpm and 37°C. Prior to SDS-PAGE, samples were heated for 10 min at 95°C.

### Functional quantitative proteomics experiments using 2nSILAC and pSILAC

For the 2nSILAC-based loss-of-function experiment, cells were differentially labeled with arginine and lysine, including a label switch. Cell were treated as indicated and mixed in equal amounts based on cell numbers. Mixed cells were washed with ddH_2_O, resuspended in 500 µl of urea buffer (8 M urea, 75 mM NaCl, 50 mM Tris-HCl pH 8.0). Cells were disrupted by bead beating using a Minilys^®^ homogenizer three times for 2 min at 4,000 rpm with at least 2 min cooling on ice between the cycles. Cell debris was removed by centrifugation for 5 min at 15,000 x g and 4°C. Protein concentrations were determined using the Bradford assay (50) before proceeding with proteolytic digestion.

For 2nSILAC-based analysis of detergent-insoluble protein aggregates, cells were differentially labeled with arginine and lysine and subjected to heat stress at 46°C for 12 min. Cells were equally mixed based on cell numbers before proceeding with biochemical enrichment of detergent-insoluble protein aggregates. Proteins were separated on SDS gels and subjected to tryptic in-gel digestion.

In pulsed SILAC experiments, cells were grown in ‘light’ SC medium and subjected to heat stress at 42°C for 30 min. Afterwards, cells were harvested by centrifugation (2 min at 1,600 x g) and resuspended in either ‘medium-heavy’ or ‘heavy’ SC medium and incubated at 30°C. Samples were taken 0, 30, 60, 90 and 120 min after heat stress and cells were mixed in equal amounts based on cell numbers. Cell lysates for MS analysis were prepared as described above.

### Preparation of samples for mass spectrometry

10 µg of protein were used for proteolytic in-solution digestion. Cysteine residues were reduced with TCEP, alkylated with iodoacetamide and proteins were digested using LysC and trypsin as previously described (22). Proteolytic in-gel digestion was performed as previously described (46). Peptides derived from in-solution or in-gel digestion were desalted using StageTips as described before with minor modifications (51). In brief, peptide mixtures were loaded onto StageTips conditioned with methanol, followed by 80% (v/v) acetonitrile (ACN)/0.5% (v/v) acetic acid and 0.5% (v/v) acetic acid. Peptides were washed twice using 0.5% (v/v) acetic acid and eluted using 80% (v/v) ACN/0.5% (v/v) acetic acid. Solvents were evaporated and peptides were resuspended in 0.1% (v/v) trifluoroacetic acid.

### Mass spectrometry

Nano-HPLC-ESI-MS/MS analyses were performed using Ultimate 3000 RSLCnano systems (Thermo Fisher Scientific, Dreieich, Germany) online coupled to a Q Exactive Plus or an Orbitrap Elite mass spectrometer. The RSLC system connected to the Q-Exactive was equipped with PepMap C18 precolumns (Thermo Scientific; length: 5 mm; inner diameter: 0.3 mm; loading flow rate: 30 µl/min) and an Acclaim PepMap analytical column (Thermo Scientific; length: 500 mm; inner diameter: 75 µm; particle size: 2 µm; packing density: 10 nm; flow rate: 0.25 µl/min). The RSLC system connected to the Orbitrap Elite was equipped with a nanoEase™ M/Z Symmetry C18, 100Å, 5µm precolums (Waters; length: 20 mm; inner diameter: 180 µm; flow rate: 5-10 µl/min) and a nanoEase™ M/Z HSS C18 T3 Col analytical column (Waters; length: 250 mm; inner diameter: 75 µm; particle size: 1.8 µm; packing density: 100 Å; flow rate: 0.3 µl/min). Peptides were washed and concentrated on precolumns and separated on the analytical column using a binary solvent consisting of solvent A (Q-Exactive Plus: 0.1% [v/v] formic acid [FA]; Orbitrap Elite: 0.1% [v/v] FA/4% [v/v] DMSO) and solvent B (Q-Exactive Plus: 86% [v/v] ACN/0.1% [v/v] FA; Orbitrap Elite: 0.1% [v/v] FA/4% [v/v] DMSO/48% [v/v] methanol/30% [v/v] ACN). Peptides analyzed on the Q Exactive Plus derived from 2nSILAC-based loss-of-function study were eluted using a gradient starting with 1% solvent B for 3 min, followed by 1-2.5% B in 12 sec, 2.5-22% B in 142 min, 22-39% B in 60 min, 39-54% B in 10 min, 54-95% B in 3 min, and 5 min at 95% B. Peptides analyzed on the Q Exactive Plus derived from the analysis of detergent-insoluble protein aggregates were eluted using a gradient starting with 1% solvent B for 3 min, 1-24% B in 70 min, 24-42% B in 23 min, 42-95% B in 2 min, and 5 min at 95% B.. Peptides analyzed on the Orbitrap Elite derived from pSILAC experiments were eluted using a gradient starting with 7% solvent B for 5 min, followed by 7-50% B in 245 min, 50-95% B in 85 min, and 5 min at 95% B.

Mass spectrometers were equipped with a Nanospray Flex ion source with DirectJunction™ adaptor (Thermo Scientific) and fused silica emitter (PicoTip, New Objective; Q-Exactive Plus) or distal coated fused silica Tips (FS360-20-10-D, New Objective, Woburn, USA; Orbitrap Elite). The MS instruments were operated in data-dependent acquisition mode. For the Q Exactive Plus, full MS scans were acquired with a mass range of *m/z* 375 – 1700, a resolution of 70,000 at *m/z* 200, a maximum automatic gain control (AGC) of 3 x 10^6^ ions, and a maximum injection time (IT) of 60 ms. The 15 most intensive peptide ions (z ≥ 2) were selected for fragmentation by higher-energy collisional dissociation (HCD). MS/MS scans were acquired with a normalized collision energy (NCE) of 28%, an AGC target of 1 x 10^5^ ions, a maximum IT of 120 ms, an isolation window of 3.0 *m/z,* a resolution of 35,000, and a dynamic exclusion (DE) time of 45 sec. For the Orbitrap Elite, full MS scans were acquired with the following parameters: mass range, *m/z* 370 – 1700; resolution, 120,000 at *m/z* 400; AGC, 1 x 10^6^ ions; maximum injection time, 200 ms. The 25 most intensive peptide ions (z ≥ 2) were selected for fragmentation by collision-induced dissociation (CID). MS/MS scans were acquired with the following parameters: NCE of 35%, activation q of 0.25, activation time of 10 ms, target value of 5,000, maximum IT of 150 ms, isolation width of 2.0 *m/z,* and DE of 45 s.

### Data Analysis

MaxQuant with the integrated search engine Andromeda was used for protein identification and quantification (version 1.5.5.1) (52, 53). MS/MS data were searched against a yeast-specific database obtained from the *Saccharomyces* Genome Database (SGD; http://www.yeastgenome.org/) and a list of common contaminants included in the MaxQuant software package. Database searches were carried out using trypsin and LysC specificity, a maximum number of two missed cleavages, and a mass tolerance of 4.5 ppm for precursor and 0.5 Da (CID data) or 20 ppm (HCD data) for fragment ions. Oxidation of methionine and acetylation of protein N-termini were considered as variable and carbamidomethylation of cysteine residues as fixed modification. Multiplicity was set to three, and Arg10/Lys8 were set as ‘heavy’, Arg6/Lys4 as ‘medium-heavy’ and Arg0/Lys as ‘light’ label. The options ‘re-quantify’ and ‘match between runs’ were enabled. Protein identification was based on peptides with a minimal length of seven amino acids and at least one unique peptide. The false discovery rate for peptide and protein identifications was 0.01. For 2nSILAC-based relative protein quantification, a minimum ratio count of one was required.

In 2nSILAC-based experiments, mean log_2_ ratios of normalized SILAC ratios were calculated and a two-sided Student’s *t*-test was performed to determine *p*-values. For outlier analysis, the Perseus software (version 1.6.14) (54) was used to determine Significance B *p*-values (52). Proteins with a *t-*test p-value and a Significance B *p-*value of ≤ 0.05 were considered as significantly changed in abundance. YeastMine (55) was used to perform GO term enrichment analyses. GO terms with a Holm-Bonferroni corrected p-value of ≤ 0.05 were considered as over- or underrepresented. Translation rates (56), half lifes (57), protein abundance (46), physio chemical properties (derived from the SGD (58)) were analyzed using R & RStudio.

In pSILAC experiments, non-normalized H/M ratios were used to assess differences in global protein synthesis between the two populations. To correct for inequalities in sample mixing ratios, iBAQ intensities of the different populations were considered and the correction factor λ was determined as follows: λ = (Σ iBAQ intensities)*_Δola1_*/(Σ iBAQ intensities)_wt_ (40). For hierarchical cluster analysis, proteins quantified at timepoints t2-t4 (i.e. 60-120 min) were considered. In case quantitative data were missing for timepoint t1, missing values were imputated on the basis of a normal distribution. A Pearson’s correlation-based distance matrix was used for the hierarchical clustering with Ward’s method.

### Laser scanning confocal microscopy

For fluorescence microscopy, 10 ml of cells from a culture in logarithmic growth phase (OD_600_ 0.5) were fixed as previously described (59), resuspended in 20 µl of PBS containing 0.1% (v/v) Triton X-100 and analyzed using a Zeiss LSM 880 (Carl Zeiss AG, Oberkochen, Germany) equipped with a W-Plan Apochromat 63x/1.0 water objective and an Airyscan detection unit. GFP was excited with a 488-nm line, and the emission was collected with a detector range of 415-735 nm. A 561-nm line was used to excite mCherry and emission was collected with a detector range of 410-696 nm. To cover the fluorescence signal of the entire cell, 14-20 image frames were recorded with a 0.17-0.21 µm distance between Z-stack frames. Airyscan images were processed using ZEISS ZEN Black software (version 2.3) and further processed with ImageJ (60) applying the same parameters to all images.

### Purification of Ola1p

Plasmid backbones for virus production were provided by the protein expression and purification facility at MPI-CBG. MBP-Ola1p and MBP-Ola1p^GFP^ were expressed and purified from insect cells using a baculovirus expression system (Hoell et al. 2011; Jarvis, 2014). After a 3-day transfection, cells were lysed with the LM10 Microfluidizer® in lysis buffer (50 mM Tris-HCl pH 7.5, 1 M KCl, 2 mM EDTA, 1 mM DTT, 0.01 U/ml benzonase and 1x EDTA-containing protease inhibitor cocktail [Roche Applied Sciences]). The lysate was cleared by centrifugation at 18,000 x g for 45 min at 4°C. The lysate was incubated with amylose resin (New England Biolabs), and MBP-Ola1p or MBP-Ola1p-monoGFP was eluted in elution buffer (50 mM Tris-HCl pH 7.5, 1 M KCl, 2 mM EDTA, 1 mM DTT, 20 mM maltose). The MBP tag was cleaved with His-tagged Precission protease overnight at 4°C. Proteins were loaded onto an Äkta Pure chromatography setup and purified by size exclusion chromatography using a HiLoad 16/600 Superdex 200 pg column (GE Life Sciences) using storage buffer (50 mM Tris-HCl pH 7.5, 500 mM KCl, 2 mM EDTA, 1 mM DTT). Purified Ola1p and Ola1p^GFP^ were concentrated to ∼ 200 or 400 µM, respectively, flash frozen and stored at −80°C.

### Calculation of cellular Ola1p concentration

The number of Ola1p molecules is at around 42,000 copies (N) per cell (average taken from the *Saccharomyces* Genome Database (55)). Excluding the nucleus, organelles and large macromolecular complexes, the average haploid cell volume (V_cell_) of *S. cerevisiae* is 32 µm^3^ (Uchida et al. 2011). We determined the cytosolic molar concentration of Ola1p using the formula C=N/N_A_*V_cell_ in which N_A_ denotes the Avogadro constant. Thus, the cytosolic molar concentration of Ola1p is in the range of 2-2.5 µM.

### *In vitro* assays for microscopy

For microscopy experiments, Ola1p and Ola1p^GFP^ were diluted to a final concentration of 2.5 µM in a 1:10 ratio of Ola1p to Ola1p^GFP^ in 20 mM PIPES or MES buffer of varying pH with a final salt concentration 100 mM KCl and 1 mM tris(2-carboxyethyl)phosphine (TCEP). For temperature experiments, protein samples were heated at the indicated temperature for 10 min in a thermoblock. Samples were imaged at room temperature with a Nikon Eclipse Ti2 widefield microscope and a Plan Apochromat VC 60x/1.2 NA water objective (Nikon Instruments, Amsterdam, Netherlands).

### Nano-differential scanning fluorimetry (nanoDSF)

NanoDSF coupled to scattering measurements were done with the Prometheus NT.48 (Nanotemper, Munich, Germany). Ola1p was diluted to the indicated concentrations in 20 mM PIPES pH 7.0 buffer with a final concentration of 100 mM KCl and 1 mM TCEP. Samples were loaded in high-sensitivity capillaries (Nanotemper, Munich, Germany) and heated with a 0.5°C/min temperature ramp from 20°C to 70°C.

### Dynamic light scattering (DLS)

DLS measurements were done using the Prometheus Panta (Nanotemper, Munich, Germany). For DLS measurements with good signal-to-noise, 10 µM or 0.5 mg/ml Ola1p in 20 mM PIPES pH 7.0 buffer with a final salt concentration of 100 mM KCl and 1 mM TCEP was used. Samples in the presence and absence of 2 mM ATP with 2 mM MgCl_2_ were loaded in high-sensitivity capillaries (Nanotemper, Munich, Germany) and heated with a 0.3°C/min temperature ramp from 20°C to 70°C with a 2 sec DLS integration time.

### 35S *in vivo* pulse labeling

Cells (10 OD_600_ units) were harvested by centrifugation (2 min at 1,600 x g and RT) and resuspended in 10 ml of SC medium without L-methionine. To deplete the cellular L-methionine pool, cells were incubated for 5 min at 30°C in a shaking waterbath. ^35^S-methionine/cysteine mixture (10mCi/ml, Hartmann Analytic) was added to a final concentration of 6 µCi/ml and 1 ml of sample was taken every minute. To block translation, cycloheximide (50 µg/ml) was added to each sample. Cells were lysed by trichloroacetic acid precipitation as previously described (61). Precipitates were pelleted by centrifugation for 10 min at 16,000 x g and 4°C and resuspended in 80 µl of SDS sample buffer. Proteins were separated on 10% Tris-Tricine gels and imaged using a Typhoon 9410 Molecular Imager (GE Healthcare Life Sciences).

### Phos-tag gel electrophoresis

Phos-tag gel electrophoresis was performed as previously described(62). In brief, 4 ml of heat-stressed or control cells were transferred into pre-warmed test tubes and boiled for 3 min to prevent changes in phosphorylation patterns. Samples were cooled on ice and cells were collected by centrifugation. Whole cell extracts were prepared as previously described (63) and samples were dissolved in sample buffer without EDTA. For analysis of Hsf1 phosphorylation, samples were analyzed on phos-tag gels (Wako Chemicals GmbH) with 6% polyacrylamide and 20 µM phos-tag reagent, and a constant current of 10 mA per gel (Bio-Rad Mini-PROTEAN) was applied. To remove phos-tag reagent and manganese, gels were incubated for 10 min in transfer buffer containing 10 mM EDTA. Afterwards, gels were washed twice in transfer buffer without EDTA. Wet-tank Western blotting was performed for 2 h at a constant current of 375 mA. Immunoblots were decorated with α-Hsf1.

### Antibodies and immunoblotting

SDS-PAGE and Western blotting were performed according to standard procedures with polyclonal rabbit antibodies raised against Ola1p, Hsp104 (Enzo Life Sciences), Hsf1(62) and Pgk1 (Thermo Fisher Scientific) and mouse monoclonal antibodies against GFP (Roche Diagnostics), Pgk1 (Thermo Fisher Scientific) Por1/2 (Thermo Fisher Scientific) and Ubiquitin (Enzo Life Sciences). For generation of the antibody against Ola1p, the open reading frame was amplified by PCR using genomic DNA of *S. cerevisiae* strain BY4741 as template. The PCR product was digested with SacI und SalI and cloned into pET-24a (Novagen), resulting in plasmid pET24a-OLA1-6xHIS. The plasmid was transformed into *Escherichia coli* BL21, resulting in isopropyl-ß-D-thiogalactopyranosid-inducible expression of HIS6-tagged Ola1p. The tagged Ola1p was purified via IMAC using an ÄKTA Pure FPLC system (GE Healthcare) equipped with a Ni-NTA column (1 ml HisTrap HP column; GE Healthcare). Polyclonal antibodies were raised against the fusion protein (Eurogentec). Immuno-reactive complexes were detected using horseradish-peroxidase coupled anti-rabbit or anti-mouse antibodies (Sigma-Aldrich/Merck, Germany) and subsequent detection of chemiluminescence signals with a ChemoCam Camera system (INTAS Science instruments GmbH, Göttingen, Germany). Adobe Photoshop CS5 (v. 12.0.4 x64) was used for contrast adjustments and cropping.

## Data availability

The mass spectrometry proteomics data have been deposited to the ProteomeXchange Consortium (65) via the PRIDE (66) partner repository.

## Supporting information

This article contains supporting information (46, 56, 57).

## Acknowledgements

We thank Bettina Knapp for support in LC-MS analyses, Henning Jessen and Nicole Steck for scientific support and discussions, and the PRIDE team for data deposition to the ProteomeXchange Consortium.

## Funding and additional information

This work was supported by the Deutsche Forschungsgemeinschaft (DFG, German Research Foundation) Project ID 403222702/SFB 1381 (to B.W., H.G.K., S.R.), 278002225/RTG 2202 (to B.W. and H.G.K), DFG grants WA 1598/5 (to BW) and KO2184/6 (to H.G.K.) as well as the Germany’s Excellence Strategy (CIBSS – EXC-2189 – Project ID 390939984 to B.W.). Work included in this study has also been performed in partial fulfillment of the requirements for the doctoral theses of S.D. at the University of Freiburg and C.D.A. at the TU Dresden

---

**Figure S1.**
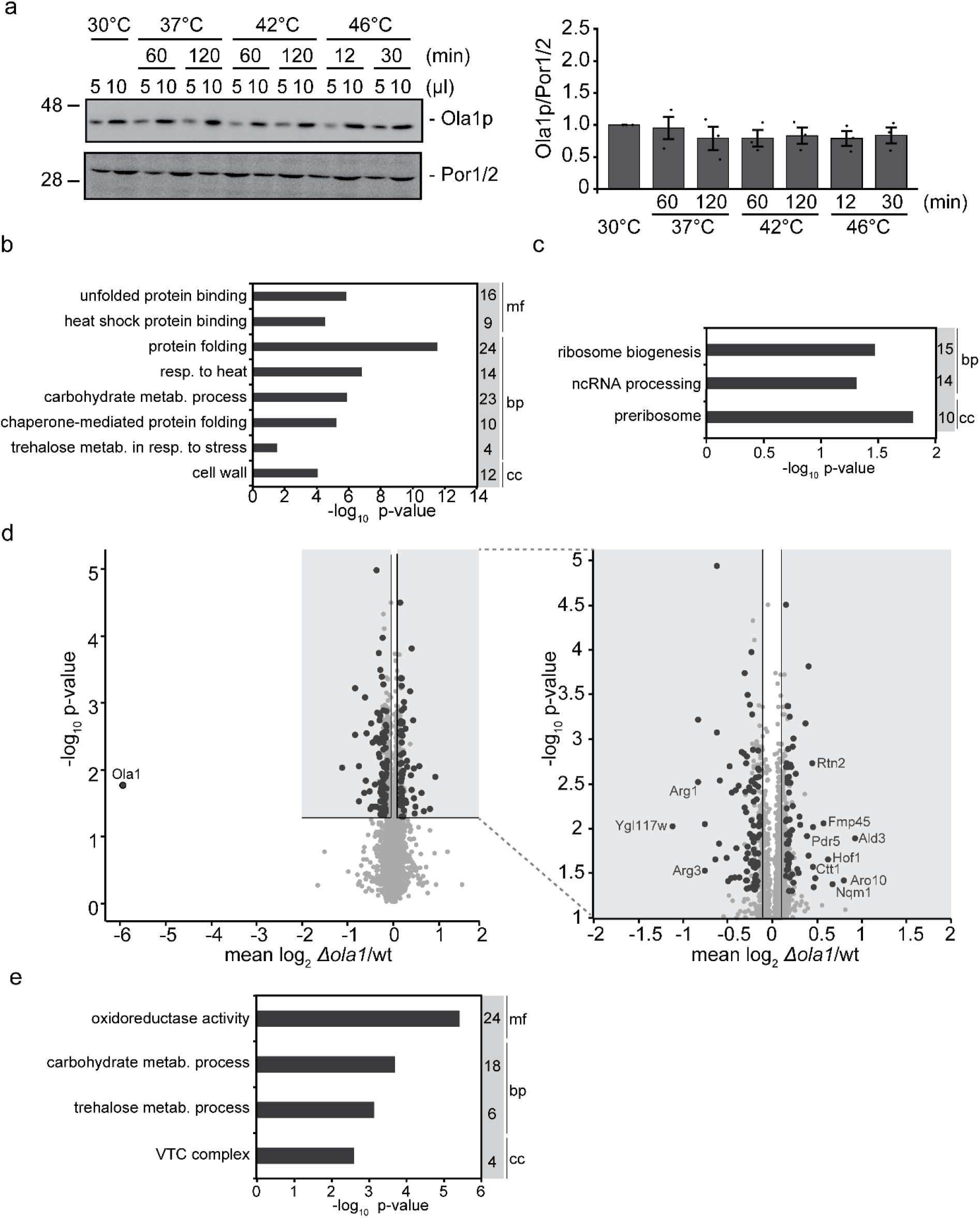
Proteomic effects of heat shock and *OLA1* deletion. a. Cells were exposed to elevated temperatures as indicated and levels of Ola1p were assessed by immunoblotting (left). Immunoblot signals for Ola1p from three independent biological replicates were quantified and normalized to the loading control Por1/2 (right). Error bars indicate SD. b, c, e. Gene Ontology term enrichment analysis of proteins significantly increased (b) or decreased (c) in abundance in heat-stressed wildtype cells (related to Figure 1b), and significantly increased in heat-stressed Δ*ola1* cells (e; related to Figure 1d). Numbers indicate the number of proteins assigned to the respective terms for the domains ‘cellular component’ (cc), ‘molecular function’ (mf), and ‘biological process’ (bp). resp., response; metab., metabolism; VTC, vacuolar transporter chaperone. d. Left, differences in the proteome of wildtype and *Δola1* cells under non-stress conditions. Larger circles indicate proteins significantly changed in abundance (i.e., p-value < 0.05 for both t-test and Significance B; n = 4). Right, Zoom-in of the shaded area of the volcano plot.

**Figure S2.**
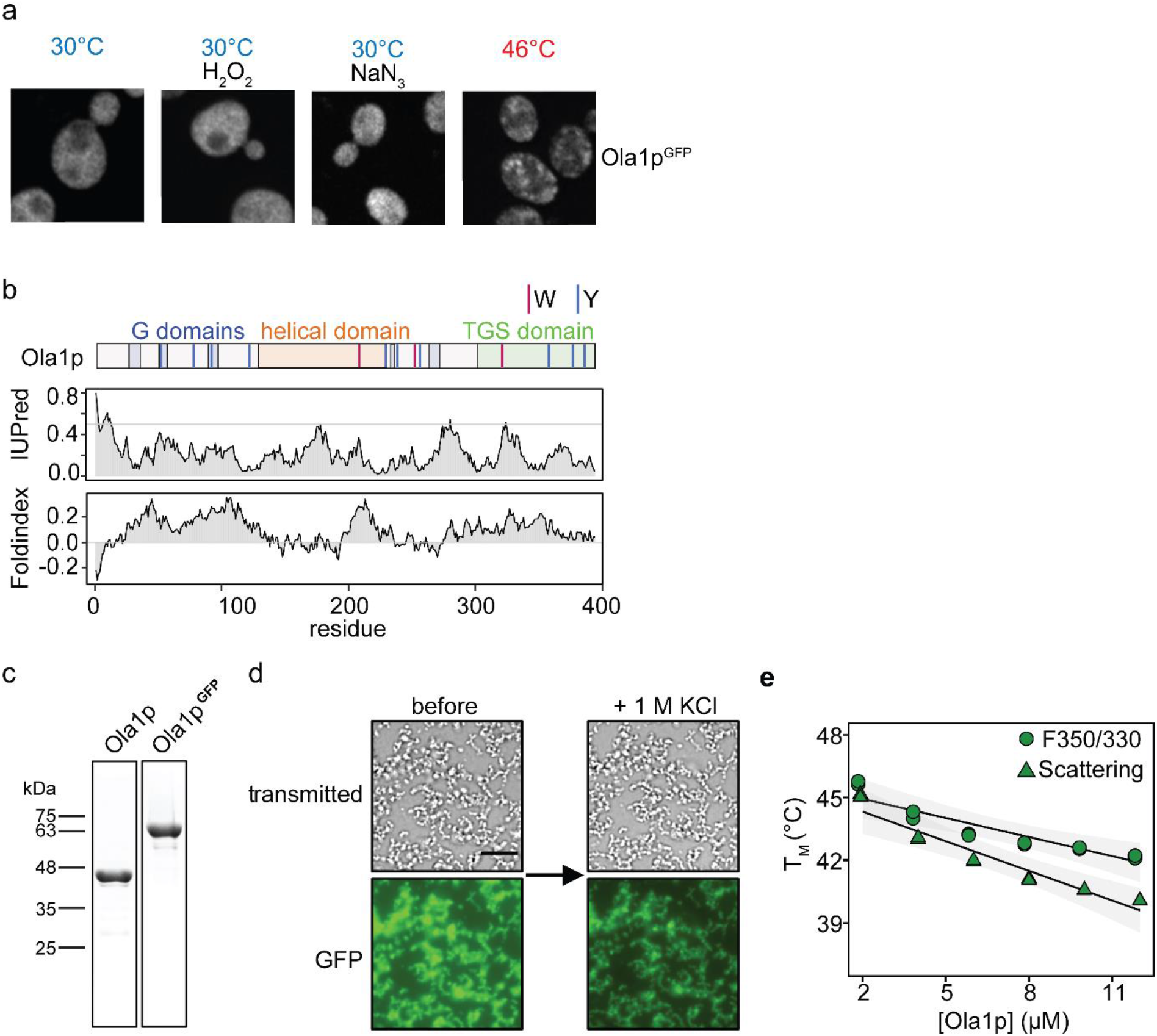
Assembly of Ola1p *in vivo* and *in vitro*. a. Subcellular localization of C-terminally tagged Ola1p^GFP^ in cells exposed to different stress conditions, analyzed by fluorescence microscopy. Stress conditions were as follows: H_2_O_2_ (1 mM, 30 min), NaN_3_ (0.5%, 30 min), and heat stress (46°C, 12 min). b. Schematic representation of the Ola1p domain structure. Tryptophan (W) and tyrosine (Y) residues are highlighted (top). Shown below are prediction plots depicting intrinsic disorder and folding of Ola1p, generated using the IUPred (68) and Foldindex (69) algorithm, respectively. c. Coomassie-stained SDS gels of purified Ola1p and Ola1p^GFP^. d. Transmitted light and fluorescent microscopy images (GFP) of 2.5 µM Ola1p heat-induced assemblies (1:10 ratio of Ola1p^GFP^ to Ola1p) in pH 7.0 buffer before and after the addition of 1 M KCl. Note that the GFP signal after addition of 1 M KCl decreases due to the dilution of the Ola1p^GFP^ signal. Scale bar, 10 µm. e. The T_M_ values of F350/330 and scattering of Ola1p are plotted as function of Ola1p concentration. Three replicates and a trend line are plotted.

**Figure S3:**
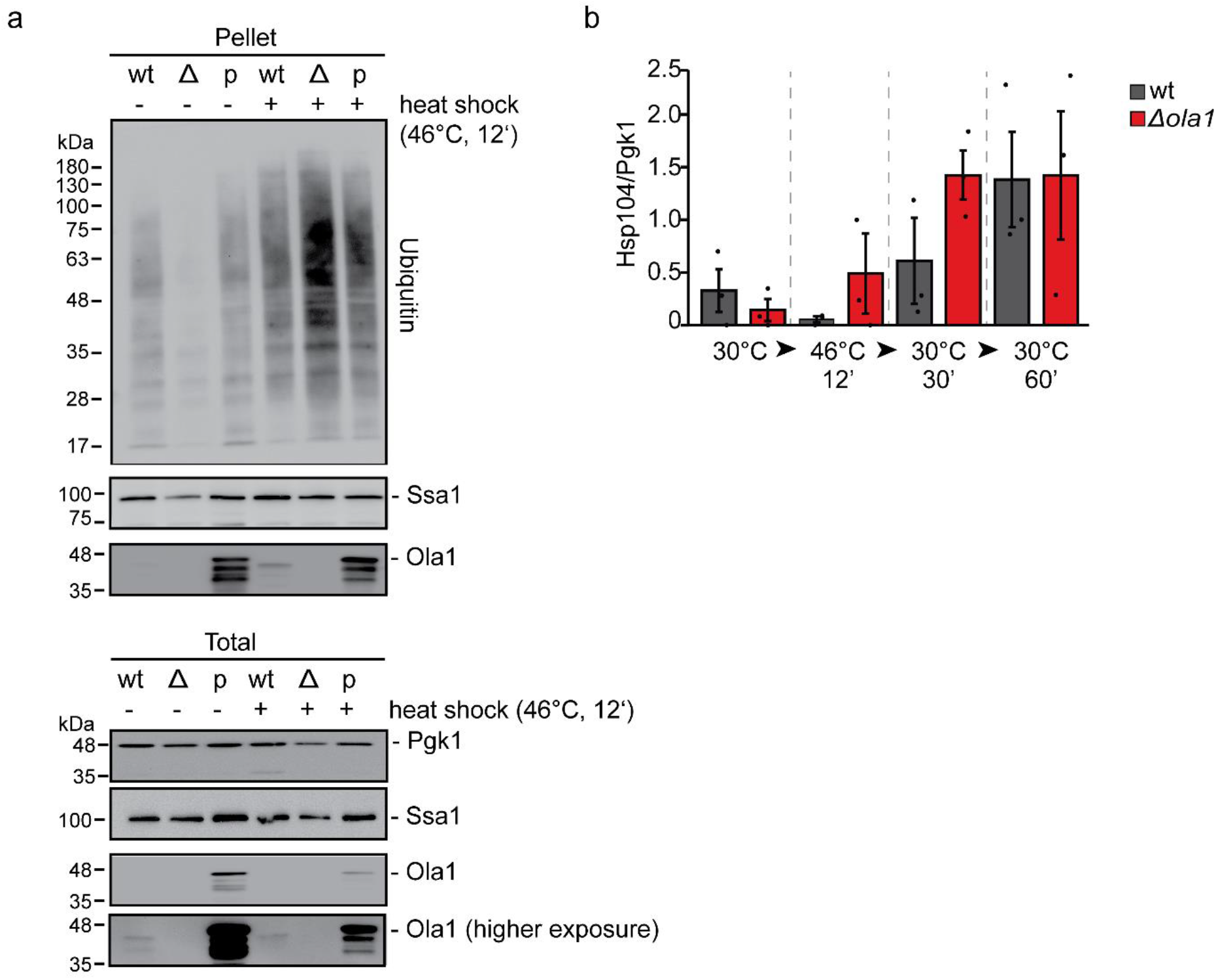
Effects of *OLA1* deletion and overexpression on detergent-resistant protein aggregates. a. Whole cell extracts (Total) and NP40-insoluble protein aggregates (Pellet) were prepared from wildtype (wt) and *Δola1* cells (Δ) as well as from cells overexpressing plasmid-encoded Ola1p (p). Fractions obtained from heat-stressed (+) and control cells (−) were analyzed by immunoblotting using antibodies directed against the indicated proteins. b. Levels of Hsp104 in protein aggregates, determined by immunoblotting as exemplarily shown in Figure 4c, were quantified (n = 3) and normalized to the signal of the loading control Pgk1 in whole cell extracts (see Figure 4d). Error bars indicate SD.

**Figure S4.**
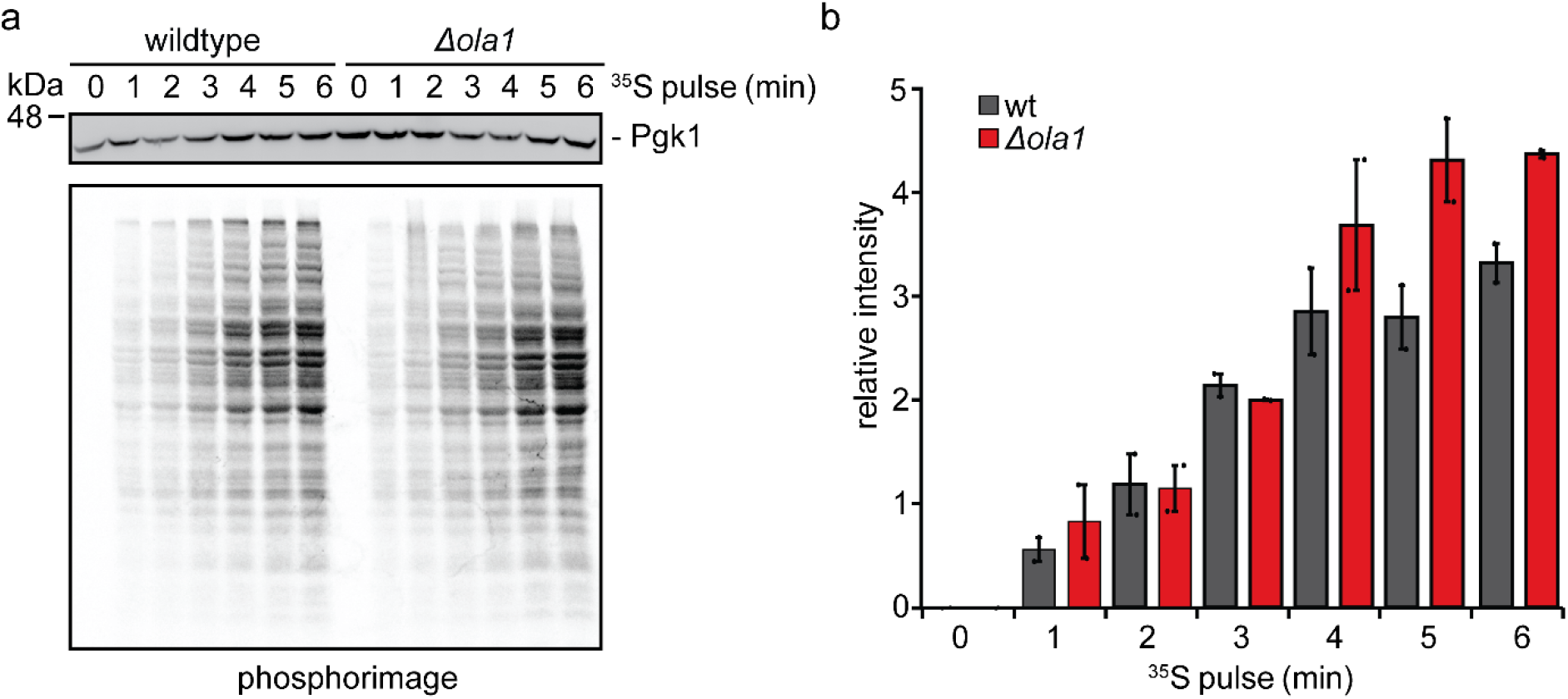
Characterization of translation reinitiation after heat shock relief. a, b. Wildtype and *Δola1* cells were cultivated at 30°C. During logarithmic growth phase, ^35^S-Cys/Met was added and the incorporation of ^35^S into newly synthesized proteins was monitored by autoradiography at the indicated timepoints (a). Autoradiographic intensities (n = 2) were quantified and normalized to the loading control Pgk1 (b). Error bars indicate SD.

**Figure S5.**
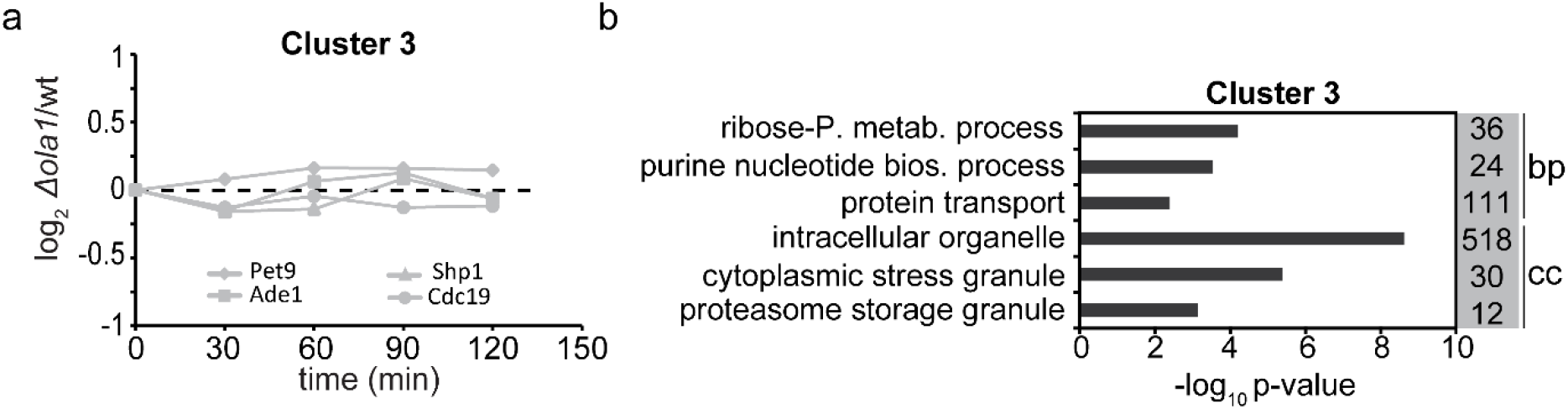
Pulsed SILAC analysis of protein synthesis during recovery from heat shock. a. Temporal profiles of relative protein synthesis of proteins representative for clusters 3 as determined by hierarchical cluster analysis described in Figure 6. b. Gene Ontology term overrepresentation analysis of cluster 3 for the domains ‘cellular component’ (cc), ‘molecular function’ (mf), and ‘biological process’ (bp). Listed are selected terms overrepresented and number of proteins assigned to each term that are present in the cluster. ribose-P. metab., ribose phosphate metabolic; bios, biosynthetic.

## Supplemental Table Legends

**Table S1. Results of the SILAC-based study of effects of heat stress on the proteomes of wildtype and Δ*ola1* cells (a) and GO term enrichment analysis (b).** Related to Figure 1 and S1.

a. Whole cell extracts of wildtype (wt) and Δ*ola1* cells treated as indicated in Figure 1a were analyzed by triple native SILAC-based quantitative MS (n = 4). Raw MS data and complete MaxQuant results files are available via ProteomeXchange with the identifier PXD025587. HS, heat stress.

b, c. GO term enrichment analysis of proteins significantly increased/decreased was performed using YeastMine (55). GO terms with a Holm-Bonferroni corrected p-value of ≤ 0.05 were considered as overrepresented. bp, biological process; cc, cellular component; mf, molecular function.

**Table S2. Results of the SILAC-based study of *OLA1* deletion on the proteome under non-stress conditions.** Related to Figure S1.

Whole cell extracts of wildtype (wt) and Δ*ola1* cells were analyzed by SILAC-based quantitative MS (n = 4). Raw MS data and complete MaxQuant results files are available via ProteomeXchange with the identifier PXD025614.

**Table S3. Results of native SILAC analysis employed to study effects of heat stress on the detergent-resistant pellet fractions of wildtype and Δ*ola1* cells.** Related to Figure 2.

a. Detergent-resistant protein aggregates, enriched via differential centrifugation from wildtype (wt) and Δ*ola1* cells, were analyzed by native SILAC-based quantitative MS (n = 3). Raw MS data and complete MaxQuant results files are available via ProteomeXchange with the identifier PXD025588.

b. GO term enrichment analysis of proteins significantly enriched in pellet fractions of heat-shocked wildtype cells was performed using YeastMine (55). GO terms with a Holm-Bonferroni corrected p-value of ≤ 0.05 were considered as overrepresented. bp, biological process; cc, cellular component; mf, molecular function.

**Table S4. Results of pulsed SILAC approach employed to study protein synthesis in wildtype and Δ*ola1* cells during recovery from heat shock.** Related to Figure 6 and S5.

a. Whole cell extracts of wildtype (wt) and Δ*ola1* cells treated as indicated in Figure 6a were analyzed by pulsed SILAC (n = 3). Raw MS data and complete MaxQuant results files are available via ProteomeXchange with the identifier PXD025673

b. GO term enrichment analysis of proteins significantly overrepresented in clusters 1-3 as determined by hierarchical clustering was performed using YeastMine (55). GO terms with a Holm-Bonferroni corrected p-value of ≤ 0.05 were considered as overrepresented. bp, biological process; cc, cellular component; mf, molecular function.

**Table S5. Results of pulsed SILAC analysis employed to study protein synthesis in wildtype and Δ*ola1* cells overexpressing Ola1p during recovery from heat shock.** Related to Figure 6g.

Whole cell extracts of wildtype (wt) and Δ*ola1* cells overexpressing Ola1p were treated as indicated in Figure 6a and analyzed by pulsed SILAC (n = 3). Raw MS data and complete MaxQuant results files are available via ProteomeXchange with the identifier PXD025615.

